# Hepatocyte Membrane Potential Regulates Serum Insulin and Insulin Sensitivity by Altering Hepatic GABA Release

**DOI:** 10.1101/475608

**Authors:** Caroline. E. Geisler, Susma. Ghimire, Chelsea. Hepler, Kendra E. Miller, Mark. R. Higgins, Jun. Yoshino, Samuel. Klein, Benjamin. J. Renquist

## Abstract

The degree of hepatic lipid accumulation in obesity correlates with the severity of hyperinsulinemia and systemic insulin resistance. Here, we propose a mechanism that explains this associative link, whereby, hepatic steatosis dysregulates glucose and insulin homeostasis. Obesity-induced hepatocellular lipid accumulation results in hepatocyte depolarization. We have established that hepatocyte depolarization depresses hepatic afferent vagal nerve firing, increases GABA release from liver slices, and causes hyperinsulinemia. Preventing hepatic GABA release or eliminating the ability of the liver to communicate to the hepatic vagal nerve ameliorates the hyperinsulinemia and insulin resistance associated with diet-induced obesity. In people with obesity hepatic expression of GABA transporters is associated with glucose infusion and disposal rates during a hyperinsulinemic euglycemic clamp. Single nucleotide polymorphisms in hepatic GABA re-uptake transporters are associated with an increased incidence of type II diabetes mellitus. Herein, we identify GABA as a novel hepatokine that is dysregulated in obesity and whose release can be manipulated to mute or exacerbate the glucoregulatory dysfunction common to obesity.

## Introduction

Non-alcoholic fatty liver disease (NAFLD) is associated with an increased risk of developing diabetes, while the degree of hepatic steatosis is directly related to the severity of systemic insulin resistance, glucose intolerance, and hyperinsulinemia ^1–3^. We hypothesized that hepatic lipid accumulation caused hyperinsulinemia and insulin resistance and aimed to identify a potential mechanism by which this could be mediated.

The literature suggests that the hepatic vagal nerve communicates with the central nervous system to affect pancreatic insulin release and peripheral tissue insulin sensitivity. The hepatic vagal afferent nerve (HVAN) regulates parasympathetic efferent nerve activity at the pancreas to alter insulin secretion ^4^. A decrease in HVAN firing frequency stimulates insulin secretion, whereas an increase in HVAN firing frequency decreases insulin secretion ^4,5^. The HVAN is also involved in regulating whole-body insulin sensitivity. Hepatic vagotomy diminishes insulin sensitivity (assessed as insulin-stimulated glucose uptake) in insulin sensitive rats, while improving insulin sensitivity and glucose tolerance in insulin resistant mice ^6,7^. Therefore, the firing frequency of the HVAN is integral to controlling insulin secretion and sensitivity.

In addition to serving as a hub of metabolism, the liver is also a key endocrine organ which produces a significant number of hepatokines that are altered by obesity, NAFLD, and exercise and signal to change metabolic function in other tissues ^8–11^. Despite the established role of the HVAN in affecting both insulin release and sensitivity, none of these hepatokines had been implicated in altering hepatic vagal afferent nerve activity. Hepatocellular lipid accumulation depolarizes hepatocytes ^12^. Because NAFLD is integral to the development of hyperinsulinemia and insulin resistance and the HVAN regulates insulin secretion and action, we tested the hypothesis that lipid-induced hepatocyte depolarization changes the release of neurotransmitters to affect firing activity of the HVAN and drive the dysregulation of systemic glucose homeostasis common in obesity.

## Results

### Hepatic Vagotomy Protects Against Diet-Induced Hyperinsulinemia

Chronic hepatic vagotomy eliminates the ability of the liver to alter vagal afferent nerve activity. However, hepatic vagotomy does not prevent otherwise basal signaling of the vagus in the central nervous system ^13^. In fact, basal signaling at the nucleus of the solitary tract (NTS) is restored within 1 month of surgery ^13^.

To investigate our hypothesis that hepatic lipid accumulation drives hyperinsulinemia and insulin resistance by altering HVAN activity, we set out to test if obesity-induced insulin dysregulation is dependent on an intact hepatic vagal nerve. We expected that hepatic vagotomy would mute obesity-induced hyperinsulinemia and insulin resistance. We performed hepatic vagotomy or sham surgery in lean mice and then provided them a 60% high fat diet (HFD; Teklad, TD 06414) for 9 weeks. The operative field is visualized in Fig. 1A, with arrow A indicating the hepatic vagal branch which was severed to hepatic vagotomize mice (Fig. 1A). Vagotomy suppressed weight gain on a HFD starting at week 6 (Fig. 1B). Hepatic vagotomy lowered serum insulin and elevated the glucose:insulin ratio at both 0 and at 9 weeks on the HFD, while decreasing serum glucose concentrations after 9 weeks of HFD feeding (Fig. 1C-1E). For the same increase in body weight during HFD feeding, the rise in serum insulin was greater in sham than vagotomized mice (Fig. 1F). Thus, the vagotomy induced protection against obesity-induced hyperinsulinemia is not due to limited weight gain on the HFD. Serum glucagon concentrations in HFD fed mice were not different between surgical treatments (Fig. 1G).

**Figure 1.**
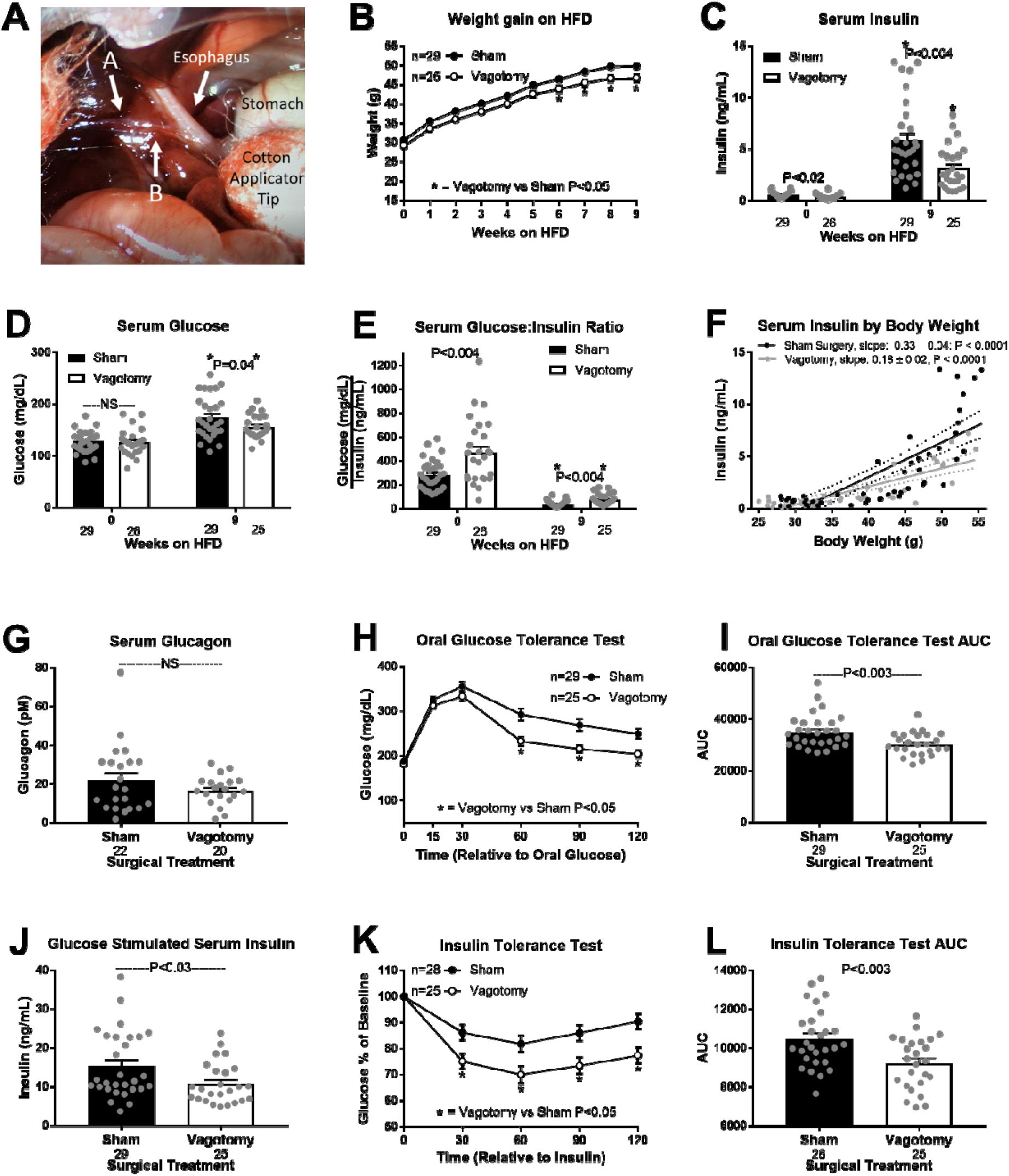
Hepatic vagotomy protects against diet-induced hyperinsulinemia. Visual operative field for hepatic vagotomy surgeries (A). Arrow A indicates the hepatic branch of the vagus which was severed to vagotomize mice. Arrow A also indicates the electrode placement to record firing activity of the hepatic vagal afferent nerve (Fig. 2F). Arrow B indicates where the hepatic vagal nerve was cut after securing the electrode to eliminate vagal efferent activity (Fig. 2F). Effects of hepatic vagotomy on high fat diet (HFD) induced weight gain (B), serum insulin (C), glucose (D), and glucose:insulin ratio (E) at 0 and 9 weeks. (C-E) * denotes significance (P < 0.05) between bars of the same color. Regression of body weight and serum insulin concentrations during HFD feeding in sham and vagotomized mice (F). Effect of hepatic vagotomy after 9 weeks of HFD feeding on serum glucagon (G), oral glucose tolerance (OGTT; H), OGTT area under the curve (AUC; I), oral glucose stimulated serum insulin (J), insulin tolerance (ITT; K), and ITT AUC (L). NS = non-significant. Number below bar denotes n per group. All data are presented as mean ± SEM.

Vagotomy improved oral glucose tolerance at 9 weeks on the HFD, while simultaneously decreasing glucose stimulated insulin concentrations (Figs. 1H-1J). Vagotomy also improved insulin sensitivity in obese mice (Figs. 1K-1L). These data support the conclusion that surgically interrupting hepatic vagal signaling attenuates the development of diet-induced hyperinsulinemia and insulin resistance.

### Hepatocyte Depolarization Depresses HVAN Firing Activity

Obesity depolarizes hepatocytes (Fig. 2D). We hypothesize that obesity-induced hepatocyte depolarization is communicated through the HVAN to dysregulate insulin secretion and action. We used the genetically-engineered, PSEM89S ligand-gated depolarizing ion channel described by Magnus, et al. (2011), to assess the effect of hepatocyte depolarization on HVAN firing activity ^14^. We intravenously delivered an adeno-associated virus serotype 8 (AAV8) encoding this PSEM89S ligand-gated depolarizing channel and green fluorescent protein (eGFP) flanked by LoxP sites to wildtype mice or mice expressing cre-recombinase driven by the albumin promoter ^15,16^. This channel will only be expressed in hepatocytes of cre-recombinase expressing mice and activated by PSEM89S ligand. We performed immunohistochemistry against GFP to confirm liver specific expression in albumin-cre expressing mice and no expression in wildtype mice (Figs. 2A-2B). No GFP expression was observed in skeletal muscle, pancreas, or adipose tissue of albumin-cre mice (Extended Data Fig. 1A). To assess the influence of hepatocyte depolarization on HVAN firing we simultaneously measured hepatocyte membrane potential and HVAN activity in the anesthetized mouse. To record HVAN firing activity, the hepatic vagal nerve was gently lifted and placed onto a hook-shaped electrode (Fig. 1A; arrow A). After the electrode placement was secured, the hepatic vagal nerve to the right of the hook near the esophagus (Fig. 1A; arrow B) was cut to eliminate efferent firing activity. Bath application of the PSEM89S ligand (30 μM) depolarized hepatocytes and decreased HVAN firing activity in albumin-cre, channel expressing mice (Figs. 2E-2F), while having no effect in wildtype mice (Figs. 2E-2F).

**Figure 2.**
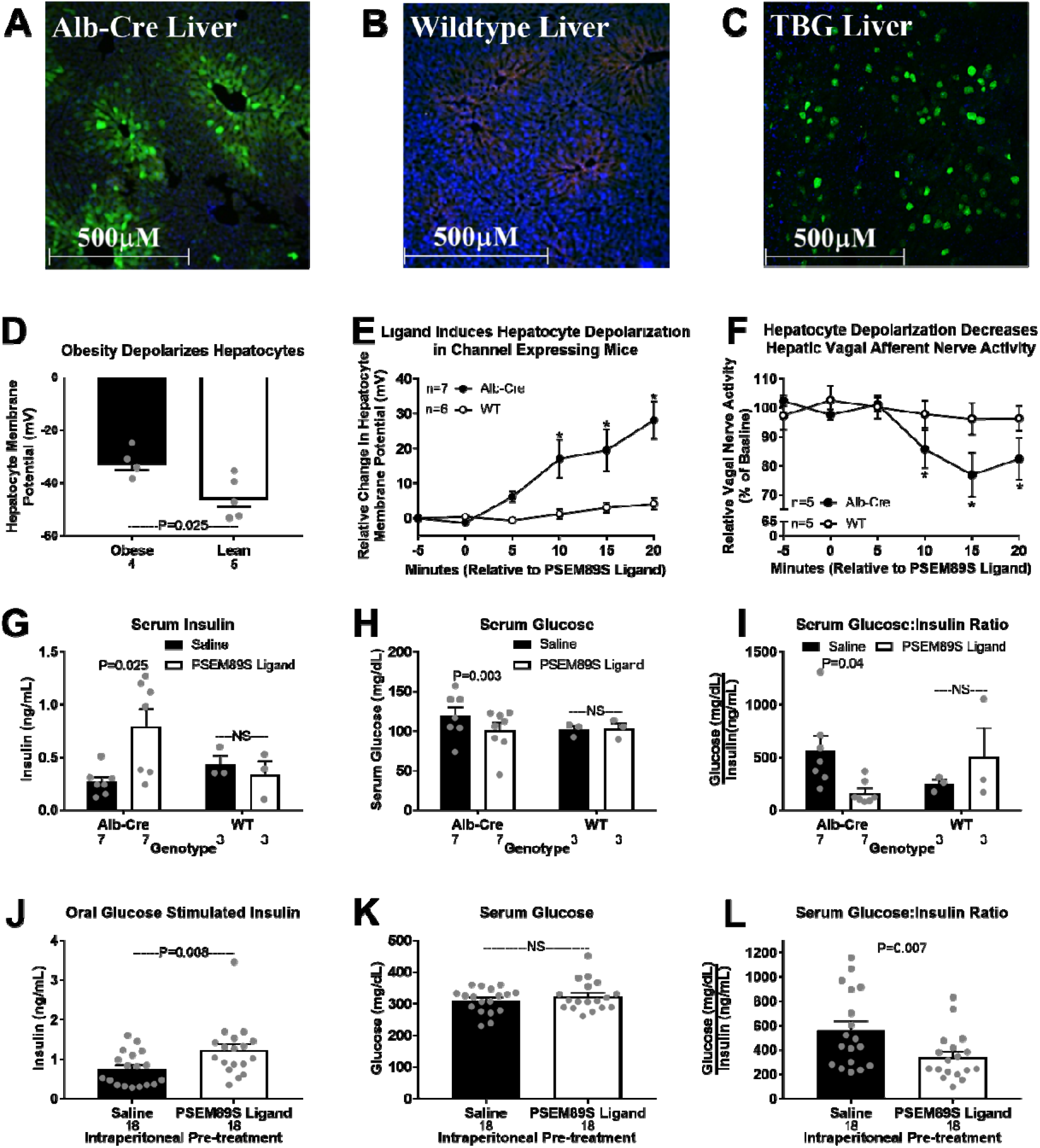
Acute hepatocyte depolarization depresses hepatic vagal afferent nerve activity and elevate serum insulin. Immunohistochemical validation of liver specific viral induced PSEM89S ligand gated depolarizing channel. (A-C; 10X magnification) Liver from an albumin-cre expressing mouse (A), and wildtype mouse (B) tail-vein injected with an AAV8 encoding the PSEM89S ligand activated depolarizing channel and green fluorescent protein (GFP) whose expression is dependent on cre-recombinase. Liver from a wildtype mouse tail-vein injected with an AAV8 encoding the PSEM89S ligand activated depolarizing channel and GFP whose expression is driven by the liver specific thyroxin binding globulin (TBG) promoter (D). Green = GFP, blue = DAPI (nucleus), and red = background fluorescence. Hepatocyte membrane potential in lean and obese mice (D). (E-I) Data from albumin-cre and wildtype mice tail-vein injected with an AAV8 encoding liver specific expression of the PSEM89S ligand activated depolarizing channel whose expression is dependent on cre-recombinase. PSEM89S ligand (30 μM) induced change in hepatocyte membrane potential (E). PSEM89S ligand induced relative change in hepatic vagal afferent nerve activity (F). Data in panel F was collected concurrently with data in panel E. Serum insulin (G), glucose (H), and glucose:insulin ratio (I) in albumin-cre and wildtype virus injected mice 15 minutes after saline or PSEM89S ligand (30 mg/kg) administration. (J-L) Data from wildtype mice tail-vein injected with an AAV8 encoding the PSEM89S ligand activated depolarizing channel whose liver specific expression is driven by the thyroxine binding globulin (TBG) promoter. Serum insulin (J), glucose (K), and glucose:insulin ratio (L) in channel expressing mice injected with either saline or PSEM89S ligand (30 mg/kg) 10 minutes prior to an oral glucose load (2.5 g/kg). Alb-Cre = albumin-cre, WT = wildtype, NS = non-significant. * denotes significance (P < 0.05) between groups within a time point. Number below bar denotes n per group. All data are presented as mean ± SEM.

### Acute Hepatic Depolarization Elevates Serum Insulin

Hepatocyte depolarization depresses HVAN activity (Figs. 2E-2F), while loss of HVAN signaling in obesity protects against the development of hyperinsulinemia (Fig. 1C). Therefore, we hypothesized that hepatocyte depolarization caused hyperinsulinemia by altering HVAN activity. To directly test this causative relationship, we intraperitoneally injected PSEM89S ligand (30 mg/kg) to induce hepatocyte depolarization and showed that this more than doubled serum insulin and decreased serum glucose concentrations in albumin-cre, channel expressing mice (Figs. 2G-2H). Accordingly, PSEM89S ligand decreased the glucose:insulin ratio in albumin-cre mice (Fig. 2I). Notably, PSEM89S ligand did not alter serum insulin, glucose, or the glucose:insulin ratio in wildtype mice (Figs. 2G-2I).

We developed a second model of hepatocyte depolarization in which liver specific expression of the same PSEM89S ligand-gated depolarizing channel and GFP was independent of cre-recombinase and instead driven by the liver specific thyroxine binding globulin (TBG) promoter ^17,18^. Wildtype mice intravenously injected with this AAV8 had liver specific GFP expression confirmed by immunohistochemistry (Fig. 2C). No GFP expression was observed in skeletal muscle, pancreas, or adipose tissue (Extended Data Fig. 1B). To further validate that the GFP positive cells in the liver are hepatocytes, we performed double immunohistochemistry for GFP and the hepatocyte specific marker, arginase-1 (Extended Data Figs. 1C-1E). To ensure stimulatory concentrations of circulating glucose, we gave an oral glucose gavage (2.5 g/kg body weight) 10 minutes following intraperitoneal PSEM89S ligand (30 mg/kg) injection. As previously observed, PSEM89S ligand administration elevated serum insulin and lowered the glucose:insulin ratio in channel expressing mice (Figs. 2J and 2L). PSEM89S ligand injection did not affect the rise in serum glucose following an oral gavage of glucose (Fig. 2K). These data establish that acute hepatocyte depolarization depresses HVAN firing activity and increases serum insulin concentrations.

### Hepatic Hyperpolarization Protects Against Diet-Induced Metabolic Dysfunction

Having established that hepatocyte depolarization increases serum insulin concentrations (Fig. 2), and that the hepatic vagus is essential for diet-induced hyperinsulinemia (Fig. 1C), we next hypothesized that hepatocyte hyperpolarization would prevent obesity-induced hyperinsulinemia. To induce a chronic hyperpolarized state, we used an AAV8 vector encoding TBG promoter driven expression of the K^+^ channel, Kir2.1, and eGFP (Fig. 3A). Although this channel is inwardly rectifying in neurons, in hepatocytes, with a resting membrane potential that ranges from −20 to −50 mV, Kir2.1 expression supports K^+^ efflux and hyperpolarization ^19^. We confirmed the hyperpolarizing effect of Kir2.1 by *in vivo* intracellular measurement of hepatocyte membrane potential before and after bath application of the Kir2.1 antagonist, Barium (Ba^2+^; 50 μM) ^19^. Ba^2+^ induced a 6.86±1.54 mV depolarization of hepatocytes in Kir2.1 expressing mice but had no effect (−0.62±1.86 mV) in control eGFP expressing mice (Fig. 3B).

**Figure 3.**
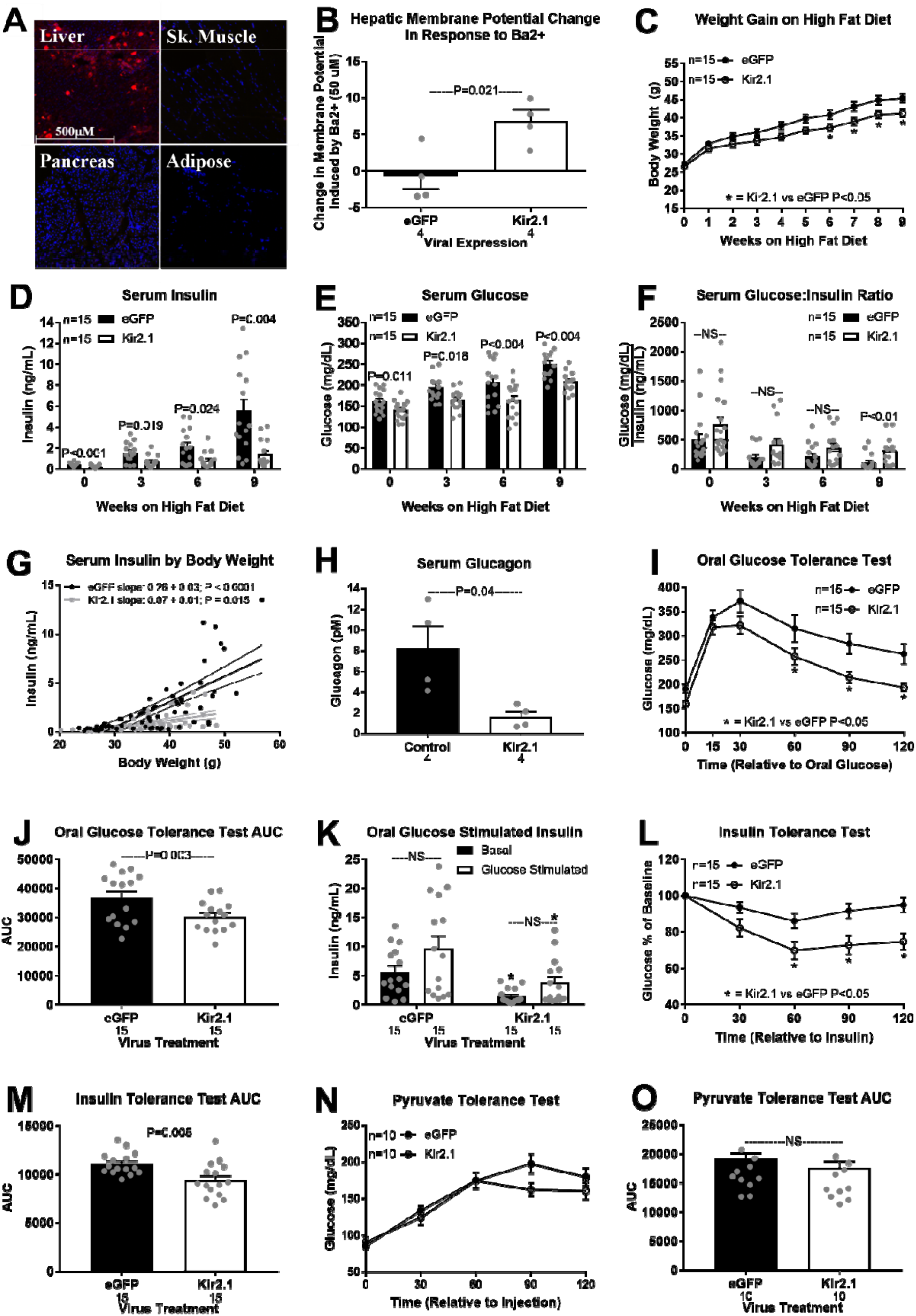
Hepatic hyperpolarization protects against diet-induced metabolic dysfunction. Liver specific expression of the Kir2.1 hyperpolarizing channel in a wildtype mouse (A; 10X magnification). Fluorescent imaging for red = tdTomato and blue = DAPI (nucleus). Barium (BaCl; 50 μM) induced change in hepatocyte membrane potential in Kir2.1 and eGFP (control) expressing mice (B). Hepatic Kir2.1 expression effect on high fat diet (HFD) induced weight gain (C), serum insulin (D), glucose (E), and glucose:insulin ratio (F) at 0, 3, 6, and 9 weeks. Regression of body weight and serum insulin concentrations during HFD feeding in Kir2.1 and eGFP mice (G). Effect of hepatic Kir2.1 expression after 9 weeks of HFD feeding on serum glucagon (H), oral glucose tolerance (OGTT; I), OGTT area under the curve (AUC; J), oral glucose stimulated serum insulin (K; * denotes significance (P < 0.05) between bars of the same color), insulin tolerance (ITT; L), ITT AUC (M), pyruvate tolerance (PTT; N), and PTT AUC (O). NS = non-significant. Number below bar denotes n per group. All data are presented as mean ± SEM.

In lean mice, hepatocyte hyperpolarization decreased basal serum insulin and glucose concentrations (Extended Data Figs. 2A-2C), improved glucose clearance (Extended Data Figs. 2D-2F) and enhanced insulin sensitivity (Extended Data Figs. 2G-2H). Kir2.1 expression did not affect gluconeogenic potential, as assessed by a pyruvate tolerance test (Extended Data Figs. 2I-2J). This establishes that hepatocyte membrane potential affects systemic glucose homeostasis in non-disease, non-obese conditions.

Kir2.1 and eGFP expressing mice were provided a HFD for 9 weeks. Kir2.1 expression depressed weight gain on a HFD, reaching significance from weeks 6-9 (Fig. 3C). As observed in lean mice, the beneficial response to hepatocyte hyperpolarization was also observed at 3 weeks on a HFD, when body weight remained similar. At 3 weeks on the HFD, Kir2.1 expression improved glucose clearance without altering glucose stimulated serum insulin (Extended Data Figs. 3A-3C). Kir2.1 expression tended to improve insulin sensitivity at 3 weeks of HFD feeding (P = 0.064; Extended Data Figs. 3D-3E). Kir2.1 expression limited the rise in serum insulin and glucose in response to 3, 6, or 9 weeks of HFD feeding, and increased the glucose:insulin ratio after 9 weeks on the HFD (Figs. 3D-3F). Although Kir2.1 expression limited HFD-induced weight gain, the same increase in body weight led to a greater increase in serum insulin concentration in eGFP control than in Kir2.1 expressing mice (Fig. 3G). Kir2.1 expression decreased serum glucagon in diet-induced obese mice (Fig. 3H). After 9 weeks on the HFD, Kir2.1 expression improved glucose tolerance and insulin sensitivity (Figs. 3I-3M), while having no effect on gluconeogenic potential from pyruvate (Figs. 3N-3O). These results suggest that hepatocyte hyperpolarization protects against the development of hyperinsulinemia, hyperglucagonemia, hyperglycemia, glucose intolerance, and insulin resistance in diet-induced obesity.

Importantly, Kir2.1 expression did not affect HFD-induced hepatic lipid accumulation (Kir2.1: 94.2±10.6 mg triglycerides/g liver versus eGFP control: 98.4±6.6 mg triglycerides/g liver; P = 0.73). The absence of hyperinsulinemia in obese Kir2.1 expressing mice despite the development of hepatic steatosis supports hepatocyte depolarization as a critical mediator in the relationship between hepatic lipid accumulation and dysregulated glucose homeostasis.

### Obesity Alters Hepatocyte Neurotransmitter Production, while Membrane Potential Affects Neurotransmitter Release

Resolving the mechanism by which hepatocyte membrane potential can alter HVAN activity and downstream glucose homeostasis is critical to understanding the link between fatty liver and insulin dysregulation. To investigate whether the liver releases neurotransmitters to affect HVAN activity, we incubated liver slices *ex vivo* and measured the release of neurotransmitters into the media (Extended Data Table 1).

Since obesity depolarizes hepatocytes (Fig. 2D), and hepatocyte depolarization decreases HVAN firing activity (Figs. 2E-2F), we hypothesized that obese livers would display either an increase in the release of inhibitory or a decrease in the release of excitatory neurotransmitters. Liver slices from obese mice released more of the inhibitory neurotransmitter GABA than liver slices from lean mice (Fig. 4A). Hepatocytes synthesize GABA via the mitochondrial enzyme GABA-Transaminase (GABA-T) ^20^. Hepatic GABA-T mRNA expression was increased in diet-induced obesity (Fig. 4B). By measuring liver triglyceride concentration in the same livers from which we had measured media GABA concentrations, we show that with an increase in liver triglyceride concentration there is increased media GABA concentration (Fig. 4C).

**Figure 4.**
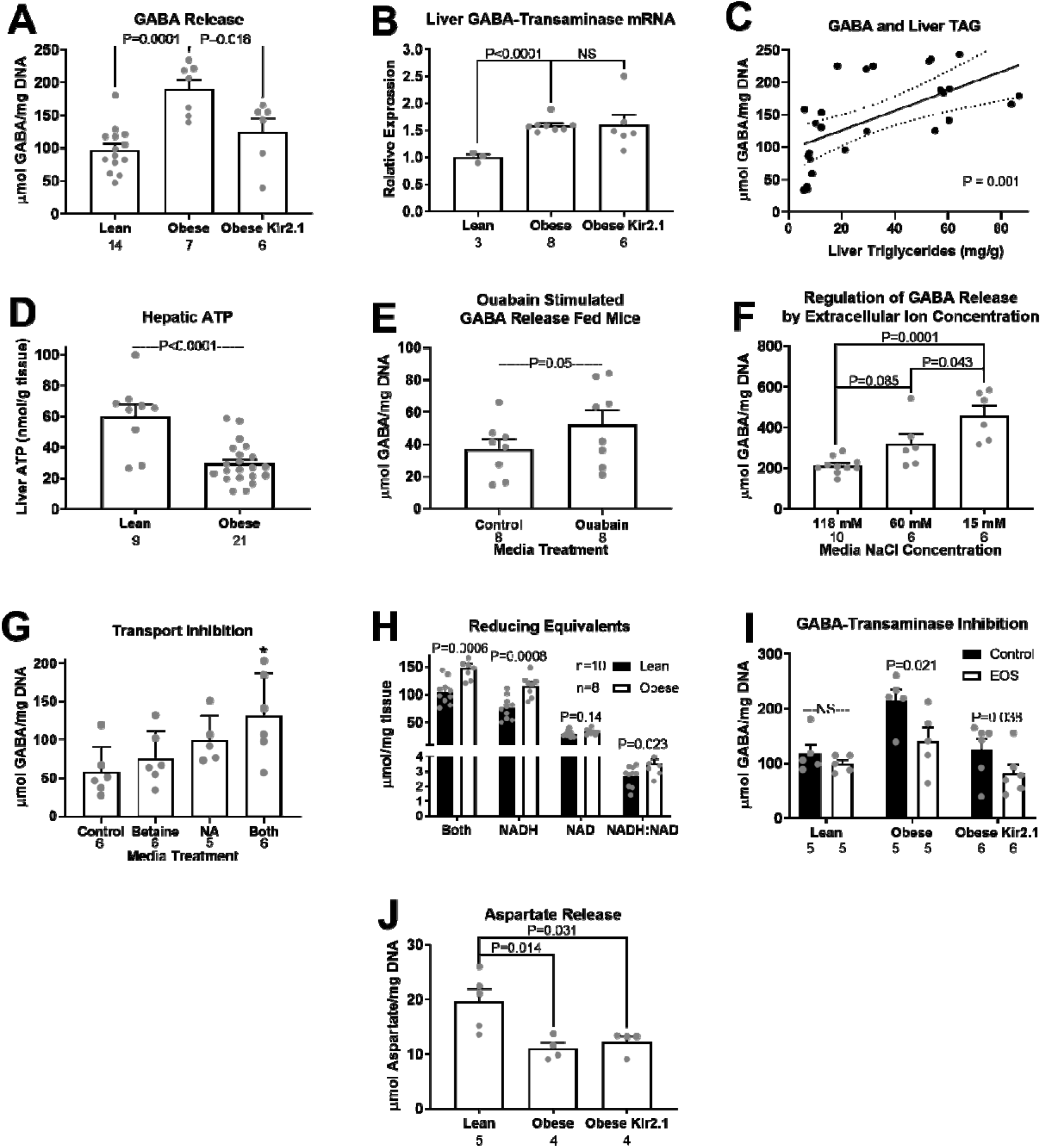
Hepatic slice GABA Release. Release of GABA (μmol/mg DNA) from hepatic slices (A), hepatic GABA-Transaminase mRNA expression (B), relationship between hepatic GABA release and liver triglyceride concentration (C), hepatic ATP concentration (nmol/g tissue; D), release of GABA in slices incubated with the Na+/K+ ATPase inhibitor, Ouabain (1 mM; E), release of GABA (μmol/mg DNA) from hepatic slices in normal (118 mM), reduced (60 mM), and low (15 mM) NaCl media (F), GABA media concentrations in slices treated with the BGT1 inhibitor (Betaine, 1 mM), the GAT2 inhibitor (Nipecotic Acid, NA, 1 mM), or both (G), reducing equivalent measures from livers of lean and obese mice (H), GABA media concentrations in response to inhibition of GABA-Transaminase (EOS, 5.3 mM) in liver slices (I), Aspartate media concentrations in lean, obese, and obese Kir2.1 expressing mice (J). *indicates difference from control (P < 0.05). NS = non-significant. Number below bar denotes n per group. All data are presented as mean ± SEM.

Diet-induced obesity decreased hepatic ATP concentrations (Fig. 4D) and lowers hepatic activity of the Na^+^/K^+^ ATPase ^21^. We propose that increased intracellular sodium ions and hepatocyte depolarization resulting from diminished Na^+^/K^+^ ATPase activity in obesity promotes GABA efflux (Fig. 5). Diet-induced obesity decreased in hepatic ATP concentrations (nmol/g tissue) to less than 50% of that seen in lean mice (Fig. 4D). Pharmacologically inhibiting Na^+^/K^+^ ATPase activity in the liver slice by bath application of Ouabain increased GABA release (μmol/mg DNA) by more than 40% (Fig. 4E). Conversely, Kir2.1 expression, which limited obesity-induced hepatocyte depolarization, decreased obesity-induced hepatocyte slice GABA release without altering the obesity-induced increase in GABA-T mRNA expression (Figs. 4A-4B). Having established a key role of membrane potential in the regulation of hepatic slice GABA release and re-uptake, we aimed to better understand hepatic GABA transport.

**Figure 5.**
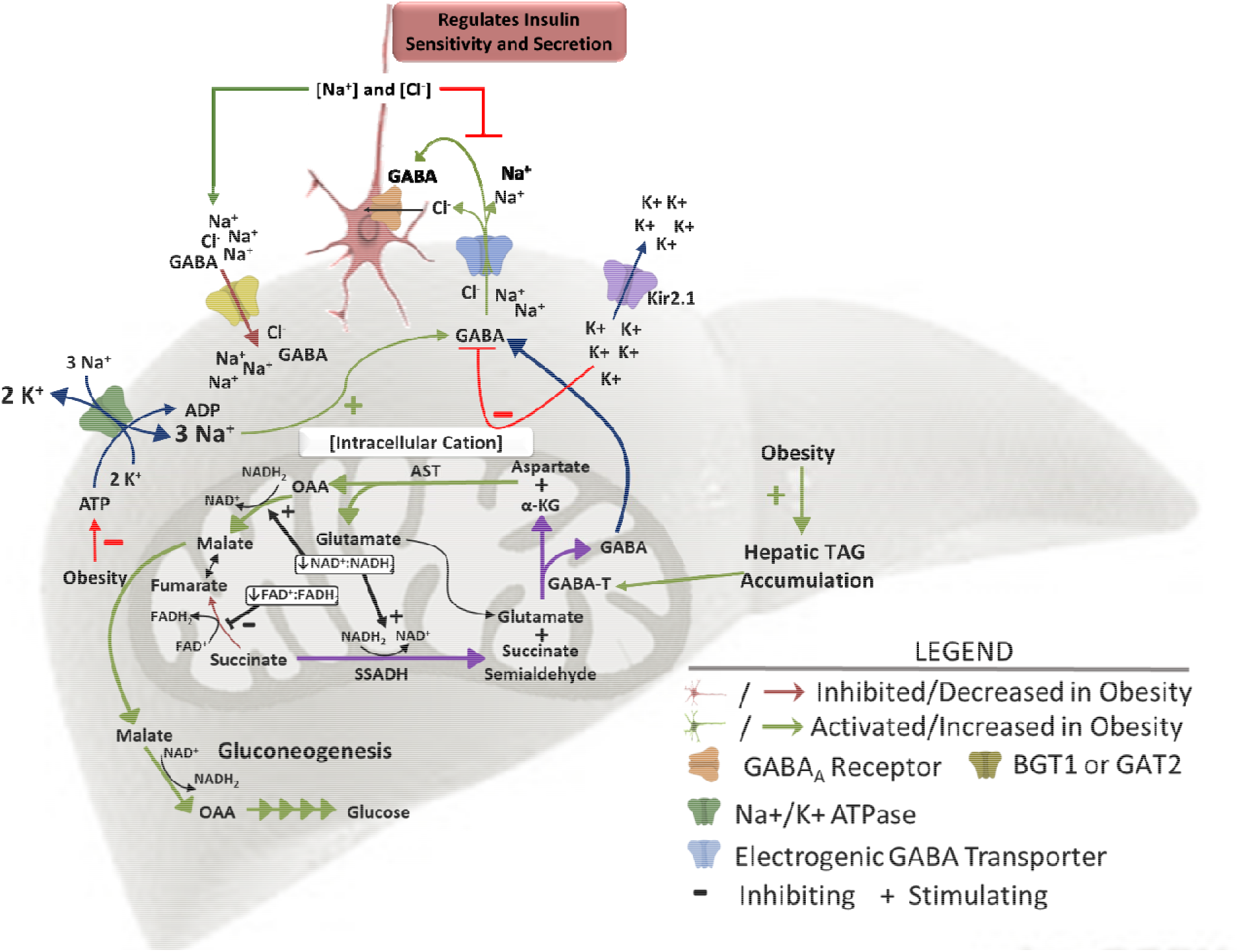
Working model of hepatic lipid accumulation induced changes in hepatic metabolism and resulting changes in hepatic vagal nerve signaling to affect insulin secretion and sensitivity. High levels of β-oxidation in the obese liver increase the mitochondrial NADH_2_:NAD^+^ and FADH_2_:FAD^+^ ratios driving succinate to succinate semialdehyde, generating substrate for GABA-Transaminase. GABA-Transaminase produces GABA and α-ketoglutarate, a substrate for aspartate aminotransferase. Increased gluconeogenic flux in obesity drives the mitochondrial export of OAA as malate. The increased GABA release is encouraged by the depolarized membrane in obesity. GABA is co-transported with 3Na^+^ and 1Cl^−^ ions, so an increase in intracellular cation concentration (hepatocyte depolarization) encourages GABA export, while a decrease in intracellular cation concentration (hepatocyte hyperpolarization) limits GABA export. Kir2.1 expression induces hepatic K^+^ efflux and hyperpolarization, inhibiting GABA export. Obesity decreases hepatic ATP concentrations, impairing activity of the Na^+^/K^+^ ATPase pump and increasing intracellular Na^+^ concentrations, driving GABA export. This mechanism explains how hepatic lipid accumulation increases hepatic GABA release. Abbreviations: OAA = oxaloacetate, AST = aspartate aminotransferase, GABA-T = GABA-Transaminase, α-KG = α-ketoglutarate, SSADH = succinate semialdehyde dehydrogenase.

Given the role of membrane potential in regulating media GABA concentrations, we proposed that liver slice GABA import and export may be mediated by ion dependent transporters. The liver expresses 4 electrogenic GABA transporters that are members of the Na^+^/Cl^−^-dependent neurotransmitter transporter (SLC6) family. These include proteins encoded for by Slc6A12 (Betaine GABA transporter 1, BGT1), Slc6A13 (GABA transporter 2, GAT2), Slc6A6 (Taurine Transporter, TauT), and Slc6A8 (Creatine transporter, CRT). BGT1 and GAT2 both co-transport 3 Na^+^, 1 Cl^−^ and GABA, moving 2 positive charges in the direction of GABA transport ^22^. TauT co-transports 2.5 Na^+^, 1 Cl^−^ and GABA, moving 1.5 positive charges in the direction of GABA transport ^23^. The CRT transporter co-transports 2 Na+, 1 Cl^−^ and GABA (or creatine) moving a single positive charge in the direction of GABA transport^24^. To establish the role of these ion dependent transporters in GABA export we showed that hepatic slice GABA release was encouraged by incubation in media with low NaCl concentrations (Fig. 4F). We also showed that incubation with the BGT1 and GAT2 inhibitors, betaine (1 mM) and nipecotic acid (NA; 1 mM), respectively, increases media GABA concentrations (Fig. 4G), establishing their redundant, but key role in hepatic GABA re-uptake (Fig. 5). As the liver is a primary sight of creatine production, it is evident that the CTR transporter can work against hepatic membrane potential to export its cargo. In fact, the reversal potential for CRT is −26 mV, well within the range of membrane potentials we report in obese mice (Fig. 2D). Additional studies will be required to fully assess the role of these electrogenic GABA transporters in GABA export. However, our studies establish the key role of membrane potential in regulating activity of these electrogenic GABA transporters results to balance GABA release and re-uptake (Fig. 5).

GABA-T is most frequently thought to be an enzyme key to GABA breakdown. However, early *in vitro* studies established that the reaction in the direction of GABA synthesis was favored with a Keq 0.04 ^25^. The reason that this reaction most frequently proceeds in the reverse direction is a lack of succinate semialdehyde (SSA), for which SSA dehydrogenase (SSADH) has a nearly 10X lower Km than GABA-T. In our model (Fig. 5), we propose that the elevated NADH:NAD ratio observed in obesity favors the production of succinate semialdehyde and ultimately GABA (Fig. 4H). To establish the key role of GABA-T activity in hepatocyte GABA release, we treated liver slices with the irreversible GABA-T inhibitor, ethanolamine-O-sulphate *ex vivo* (EOS; 5.3 mM). EOS decreased GABA export from obese control and obese Kir2.1 expressing liver slices, but not liver slices from lean mice (Fig. 4I). This supports the hypothesis that GABA production is elevated in obesity and that GABA-T mediated synthesis of GABA is not impaired by Kir2.1 expression. Hepatocytes from obese mice also released less of the excitatory neurotransmitter, aspartate, than hepatocytes from lean mice (Fig. 4J). Our model proposes that this decrease in aspartate is a direct effect of the GABA-T stimulated α-ketoglutarate synthesis and transamination to produce oxaloacetate to support gluconeogenic flux (Fig. 5). There was no effect of Kir2.1 expression on the obesity-induced decrease in aspartate release from liver slices (Fig. 4J). As hepatic Kir2.1 expression is able to prevent the hyperinsulinemia and insulin resistance in obesity (Figs. 3D and 3L) but does not affect the decrease in hepatic slice aspartate release, this proposes that a decreased excitatory signal at the HVAN in obesity is not responsible for the development of insulin resistance.

Vagal afferent innervation in the liver has previously been identified using the vagal sensory immunohistochemical marker calretinin ^26,27^. Calcitonin gene-related peptide (CGRP) has also been proposed as a marker of hepatic vagal afferent innervation and hepatic CGRP staining is eliminated by capsaicin treatment ^28^ and substantially reduced following bilateral vagotomy ^29^. We confirmed the presence of both calretinin and CGRP positive innervation in the mouse liver and established the presence of GABA_A_ receptors on calretinin and CGRP immunoreactive neurons (Figs. 6A-6D and Extended Data Fig. 4). Consistent with previous reports ^28,29^, the strongest degree of vagal afferent staining was evident in periportal areas while immunoreactive fibers penetrate into the liver parenchyma (Extended Data Figs. 4A and 4D). We propose that elevated hepatic GABA release activates GABA_A_ receptors causing chloride influx into vagal afferents explaining the decrease in HVAN activity in response to hepatocyte depolarization.

**Figure 6.**
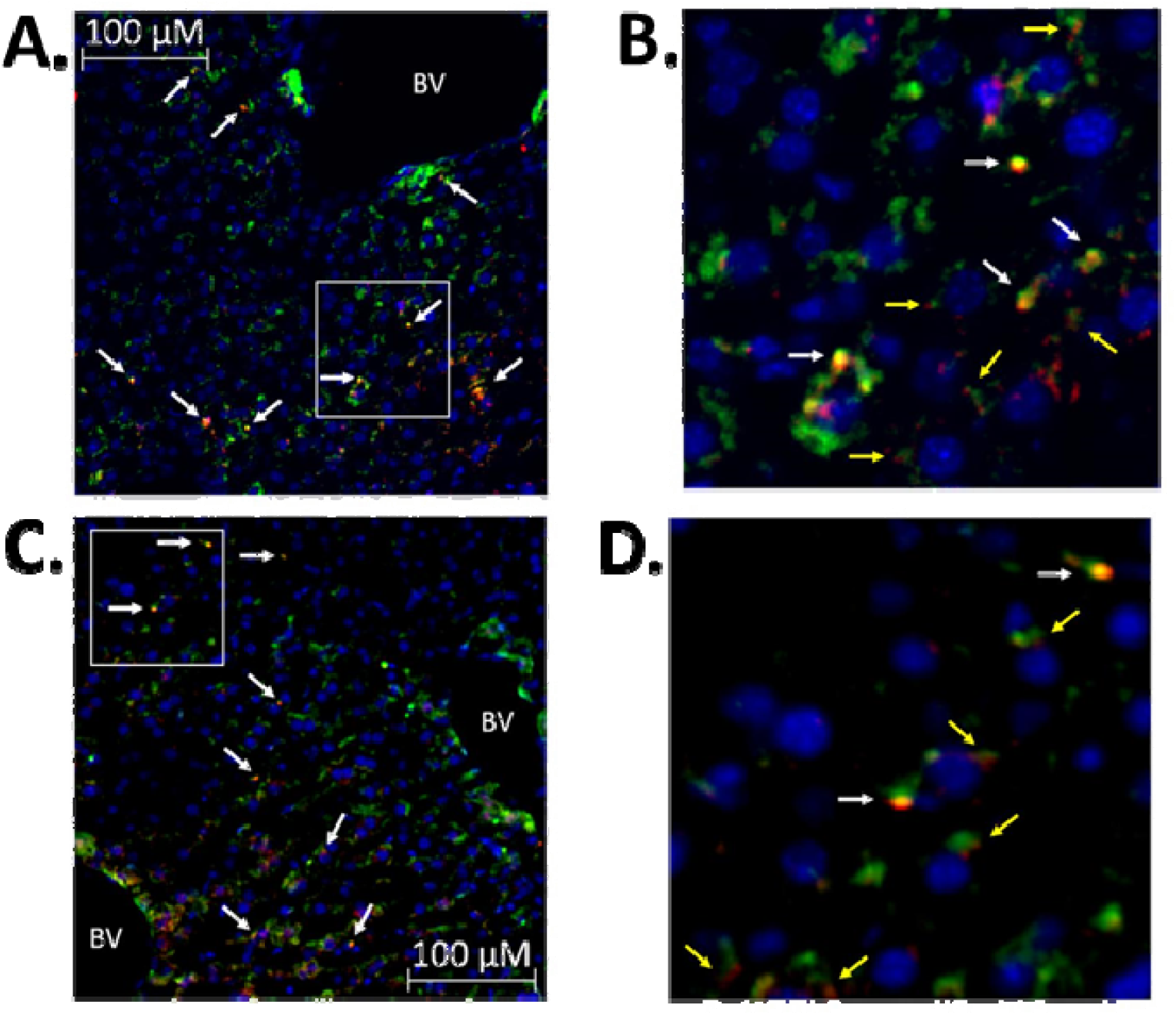
Immunohistochemical evidence of GABA_A_ receptor expressing vagal afferent innervation in the liver. Double labeling for the vagal afferent marker calretinin (green) and GABA_A_ receptors (red, A; arrows indicate co-staining). Enlarged view of area within the white box in panel A (B; white arrows indicate co-staining, yellow arrows indicate GABA_A_ positive staining immediately adjacent to calretinin positive fibers). Immunohistochemical staining for the alternative vagal afferent marker calcitonin gene-related peptide (CGRP, green) and GABA_A_ receptors (red, D; arrows indicate co-staining). Enlarged view of area within the white box in panel C (D; white arrows indicate co-staining, yellow arrows indicate GABA_A_ positive staining immediately adjacent to CGRP positive fibers). Blue = DAPI (nucleus). Image at 10X magnification. BV = blood vessel.

### Associations Between Hepatic GABA System and Glucoregulatory Markers in Humans with Obesity

To understand the potential clinical relevance of these findings, we assessed the hepatic mRNA expression of GABA transporters (*SLC6A6,* encodes for taurine transporter, TauT; *SLC6A8;* encodes for the creatine transporter, CRT; S*LC6A12;* encodes for the Betaine-GABA Transporter 1, BGT1, and S*LC6A13*; encodes for GABA Transporter 2, GAT2) in 19 people with obesity (age 45 ± 3 yrs old, 2 men and 17 women) who were carefully characterized by measuring intrahepatic triglyceride (IHTG) content using magnetic resonance imaging (MRI) and insulin sensitivity using the hyperinsulinemic-euglycemic clamp procedure (HECP) in conjunction with stable isotopically labeled glucose tracer infusion. The subjects had a wide range in IHTG content, plasma insulin concentration and measures of insulin sensitivity (Extended Data Table 2). Our multivariate regression shown that IHTG (%) was negatively associated with both glucose infusion rate during a clamp (Fig. 7A) and the percent increase in glucose rate of disposal from the basal state to the hyperinsulinemic clamp (Fig. 7B and Extended Data Table 3). Similarly, we found that hepatic S*LC*6A6 (Tau-T) and S*LC*6A8 (CRT) mRNA expression were negatively related to glucose infusion rate and insulin-induced percent increase in glucose disposal (Figs. 7A-7B). Finally, we observed that hepatic S*LC*6A12 and S*LC*6A13 expression were positively related to glucose infusion rate and insulin stimulated enhancement of glucose disposal (Figs. 7A-7B). In turn, we hypothesize that BGT1 and GAT2 are primarily acting as GABA re-uptake transporters and that Tau-T and CRT are acting to export GABA. This hypothesized role of BGT1 and GAT2 in hepatic GABA re-uptake is supported by our explant data (Fig 4G). Of note, the summed expression of *SLC6A6* and *SLC6A8* was positively associated with IHTG, suggesting that their expression increases with steatosis (P = 0.04). Again, with a reversal potential of −26 mV CRT is most likely to be a membrane potential sensitive GABA exporter (Fig. 2D).

**Figure 7.**
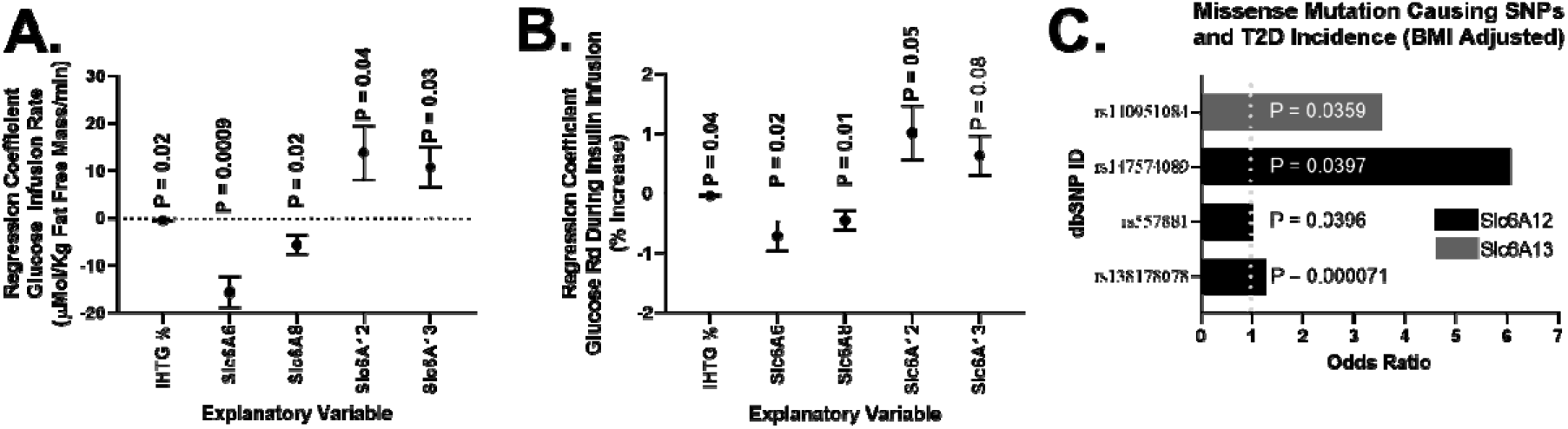
Associations between hepatic GABA system and glucoregulatory markers in obese humans. Multivariate regressions including intrahepatic triglyceride % (IHTG%) and the mRNA for the hepati GABA transporters (Slc6A6, Slc6A8, Slc6A12, and Scl6A12) as explanatory variables for variations in glucose infusion rate during a hyperinsulinemic euglycemic clamp (μMol/kg fat free mass/min; A), and the glucose disposal rate calculated during a hyperinsulinemic-euglycemic clamp (Glucose Rd, % increase; B). mRNA (FPKM; Fragments Per Kilobase of transcript per Million mapped reads) was quantified by RNA-Seq from liver tissue. (C) Single nucleotide polymorphisms (SNPs) that cause missense mutations in Slc6A12 or Slc6A13 are associated with an increased incidence of type 2 diabetes (T2D) adjusted for body mass index (BMI). Regression data are presented as mean ± SEM.

We used the Accelerating Medicines Partnership Type 2 Diabetes Knowledge Portal to understand the effect of missense mutations in genes encoding the hepatic GABA transporters (*SLC6A6*, *SLC6A8*, *SLC6A12*, and *SLC6A13*). Missense mutations in the *SLC6A12* and *SLC6A13* genes are significantly associated with an increased incidence of T2D (Fig. 7C; Extended Data Table 4). Of note a missense mutation inducing a pre-mature stop codon in the *SLC6A12* increases the odds ratio for T2D 15.8 times, establishing its key role in limiting the incidence of diabetes. Although, not specific to the liver, this GWAS data does support the hypothesis that GABA transporters encoded by *SLC6A12* and *SLC6A13* act to preventing the development of T2D.

## Discussion

We report novel neurotransmitter signaling by which hepatic steatosis may induce systemic insulin dysregulation, while establishing that hepatocyte GABA release is regulated by hepatocyte membrane potential. Using surgical and viral mouse models we have established the key role of hepatic GABA production and release and hepatic vagal nerve signaling in the dysregulation of glucose homeostasis in obesity.

We propose a new model that links hepatic lipid accumulation to HVAN activity and the development of hyperinsulinemia and insulin resistance (Fig. 5). Hepatic lipid accumulation increases flux through gluconeogenesis and increases the hepatic FADH:FAD and NADH:NAD ratio (Fig. 4H). The altered hepatic redox state inhibits the conversion of succinate to fumarate in the TCA cycle and instead drives succinate to succinate semialdehyde. Succinate semialdehyde serves as substrate for GABA-T mediated GABA production. Together with aspartate, the α-ketoglutarate formed by GABA-T produces oxaloacetate and glutamate, which feeds back into the GABA-T catalyzed reaction. The demand for gluconeogenic substrate and the high NADH:NAD ratio drives the carbons in oxaloacetate to malate and through gluconeogenesis. This gluconeogenic drive increases aspartate metabolism, explaining the decreased aspartate release in liver slices from obese mice (Fig. 4J). Thus, gluconeogenic flux and a more reduced mitochondrial redox state direct the flow of intermediate molecules in obesity resulting in elevated hepatic GABA production and increased aspartate utilization.

The ion dependence of GABA transport makes hepatocyte GABA export sensitive to changes in membrane potential. Since GABA transporters are sodium co-transporters, an increase in intracellular sodium ions and hepatocyte depolarization increases GABA export (Fig. 4E). Obesity decreases hepatic ATP content (Fig. 4D) and lowers Na^+^/K^+^ ATPase activity ^21^, providing a mechanism by which obesity depolarizes hepatocytes ^12^ (Fig. 2D) and encourages GABA export (Fig. 4A). In fact, type II diabetics have lower hepatic ATP concentrations, and both peripheral and hepatic insulin sensitivity is significantly correlated with liver ATP concentrations ^30,31^.

Our model proposes that hepatic lipid accumulation ultimately increases hepatic GABA signaling through two separate mechanisms. First, hepatic GABA production is stimulated as a result of increased GABA-T expression (Fig. 4B) and gluconeogenic flux (Fig 5), and second, hepatic GABA release is stimulated by hepatocyte depolarization (Fig. 4E). This model provides rationale to explain why gluconeogenesis and hepatocyte depolarization are essential to the development of insulin resistance and hyperinsulinemia ^32^.

This research highlights the role of hepatocyte depolarization in diet-induced metabolic dysfunction. Under physiologic conditions, hepatocyte membrane potential is closely regulated by insulin and glucagon. Acutely, insulin depolarizes while glucagon hyperpolarizes hepatocytes ^33,34^. The hyperpolarizing effect of glucagon is proposed to be mediated by cAMP and is dependent on the Na^+^/K^+^ ATPase ^33^. Accordingly, cAMP or glucagon counteracts insulin stimulated hepatocyte depolarization ^35,36^ High fat diet feeding decreases hepatic cAMP content in mice ^37^. The decrease in hepatic cAMP along with diminished ATP may contribute to obesity-induced hepatocyte depolarization. Interestingly, a loss of hepatocyte membrane polarity has been implicated in the pathology of other disease states. Hepatocellular carcinoma is characterized by hepatocyte depolarization and increased GABAergic signaling, while increasing hepatocyte polarization protects against tumor proliferation ^38^. Thus, the regulation of hepatocyte membrane potential in healthy and disease states is critically tied to cellular and metabolic function.

Physiological concentrations of insulin and glucagon induce a 5-7 mV change in hepatocyte membrane potential ^33,34^. This is comparable to the 6.86 ± 1.54 mV hyperpolarization induced by Kir2.1 expression (Fig. 3B). Admittedly, the PSEM89S ligand maximally depolarized hepatocytes by 28 ± 5.4 mV (Fig. 2E), which exceeds the depolarization observed in obesity (13 ± 4.7 mV; Fig. 2D), and likely represents a supraphysiological response. However, at 10 minutes after PSEM89S ligand administration, when HVAN activity was first significantly depressed, hepatocyte depolarization was 17.0 ± 5.4 mV, representing a more physiological change in membrane potential.

Although the idea that liver derived signals communicate to the central nervous system via the HVAN is well established in the literature ^6,39,40^, the degree hepatocyte vagal afferent innervation has remained controversial ^26^. Here, we provide evidence of vagal sensory innervation in close proximity to hepatocytes and have established the presence of GABA_A_ receptors on both calretinin and CGRP immunoreactive neurons in the liver (Figs. 6A-D) Once exported, hepatic GABA can act at GABA_A_ receptors on vagal afferents to induce chloride influx and decrease firing rate ^41^ (Fig. 5), providing a connection between hepatic lipid accumulation and decreased HVAN activity.

We appreciate that it is counterintuitive that a decrease in HVAN activity causes hyperinsulinemia (Figs. 2F, 2G, and 2J), yet surgical elimination of the HVAN limits obesity-induced hyperinsulinemia (Fig. 1C). However, surgical vagotomy preserves activity of vagal afferent axons that terminate in the hindbrain. In fact, after vagotomy NTS terminating axons show normal spontaneous activity, normal signaling from the nodose ganglion to the NTS, and evoke normal postsynaptic excitatory currents in the NTS when electrically stimulated ^13^. Re-innervation of target tissues caudal to the severed vagus is minimal out to 45 weeks post-vagotomy ^42^. Therefore, hepatic vagotomy eliminates the dysfunctional hepatocyte-vagal signaling in obesity, while preserving signaling from the HVAN above the surgical resection. Although hepatic vagotomy has been used extensively to investigate liver vagal denervation, several limitations must be acknowledged when interpreting this model ^43,44^. We cannot exclude the possibility that the improvements in glucose homeostasis in response to hepatic vagotomy (Fig. 1) are a result of interrupting non-vagal afferent signaling or partial loss of vagal efferent pancreatic innervation ^43,44^.

Studies manipulating the hepatic vagal nerve propose that afferent parasympathetic signals from the liver may affect pancreatic insulin release. Activity of the HVAN is inversely related to parasympathetic efferent nerve activity at the pancreas, which stimulates insulin release ^4,5^. Thus, portal glucose inhibits HVAN activity and increases pancreatic parasympathetic outflow to stimulate β-cell muscarinic 3 receptor (M3R) signaling and insulin release ^4,45^. Accordingly, vagotomy reduces glucose stimulated insulin secretion and basal hyperinsulinemia in obese rats by reducing cholinergic action on β-cells ^4,46^. Furthermore, cholinergic blockade decreases basal serum insulin concentrations in obese but not lean mice, suggesting that elevated basal pancreatic parasympathetic efferent tone underlies obesity-induced hyperinsulinemia ^47^.

The HVAN also regulates insulin sensitivity ^48^. Hepatic vagotomy acutely reduces insulin sensitivity in lean rats, decreasing skeletal muscle glucose clearance by 45% ^7^. In contrast, chronic hepatic vagotomy improves insulin sensitivity and glucose clearance in insulin resistant mice ^6^ (Fig. 1L). Portal glucose delivery decreases HVAN firing activity and skeletal muscle glucose clearance ^49,50^. We hypothesize that hepatic GABA production in obesity decreases HVAN activity to limit muscle glucose clearance and drive peripheral insulin resistance.

Herein, we have identified enzymes involved in GABA production and transporters involved in hepatic GABA re-uptake and release as novel therapeutic targets for correcting the inherent metabolic disturbances in T2D..

### Extended Data Titles and Legends

**Extended Data Figure 1.**
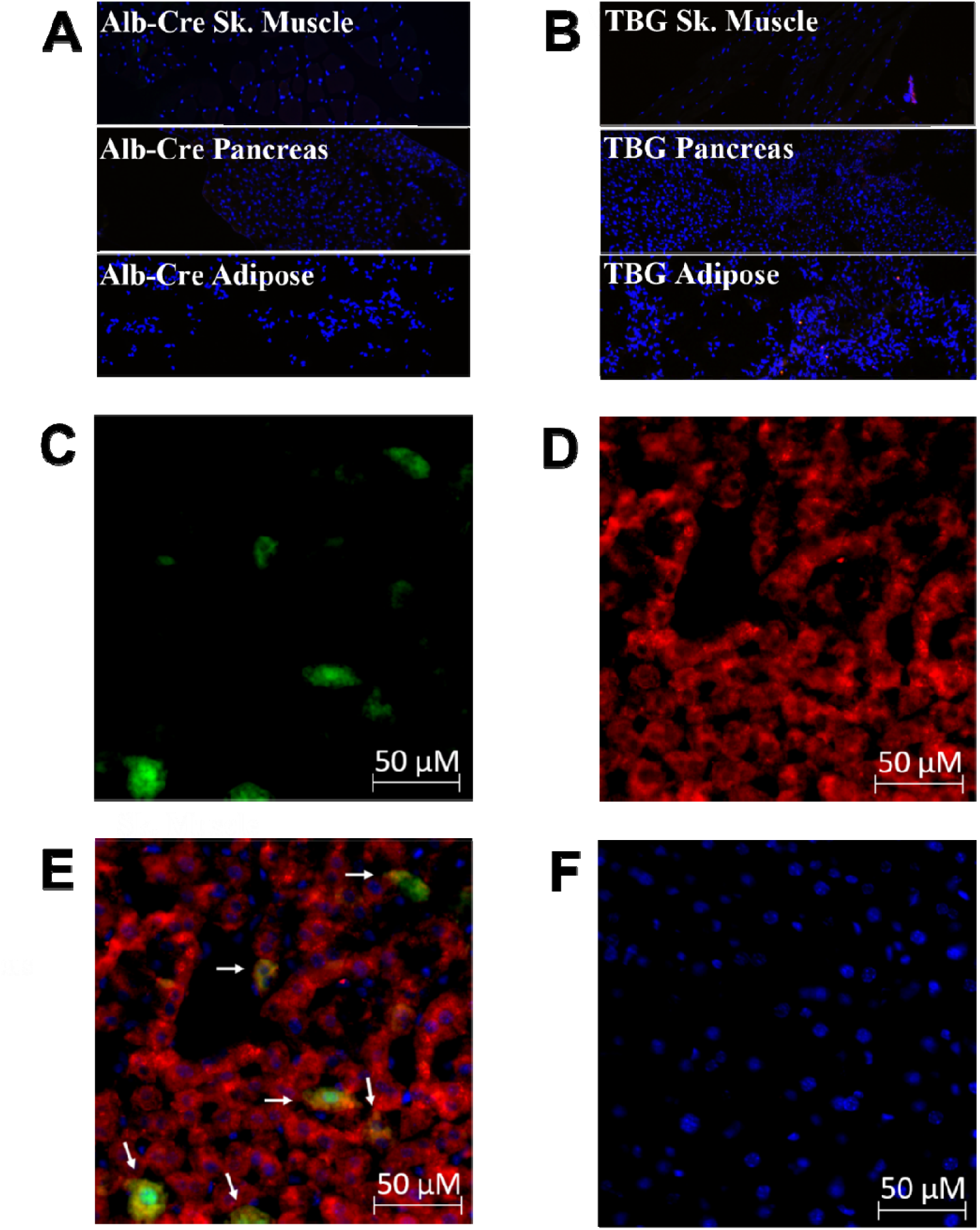
Immunohistochemical validation of liver specific viral induced PSEM89S ligand gated depolarizing channel (A-B; 10X magnification). Skeletal muscle (Sk. Muscle), pancreas, and adipose tissue from an albumin-cre expressing mouse tail-vein injected with an AAV8 encoding for the PSEM89S ligand activated depolarizing channel and green fluorescent protein (GFP) whose expression i dependent on cre-recombinase (A). Skeletal muscle (Sk. Muscle), pancreas, and adipose from a wildtype mouse tail-vein injected with an AAV8 encoding the PSEM89S ligand activated depolarizing channel and GFP whose liver specific expression is driven by the thyroxine binding globulin (TBG) promoter (B). GFP positive cells in the liver of a wildtype mouse tail-vein injected with the TBG virus co-stain with arginase-1 (C-E; 20X magnification). Staining for GFP (C), the hepatocyte specific marker arginase-1 (D), and double labeling of GFP and arginase-1 (E; arrows indicate co-staining). No primary control imaged at the same settings as panel E (F). Green = GFP, red = arginase-1, blue = DAPI (nucleus).

**Extended Data Figure 2.**
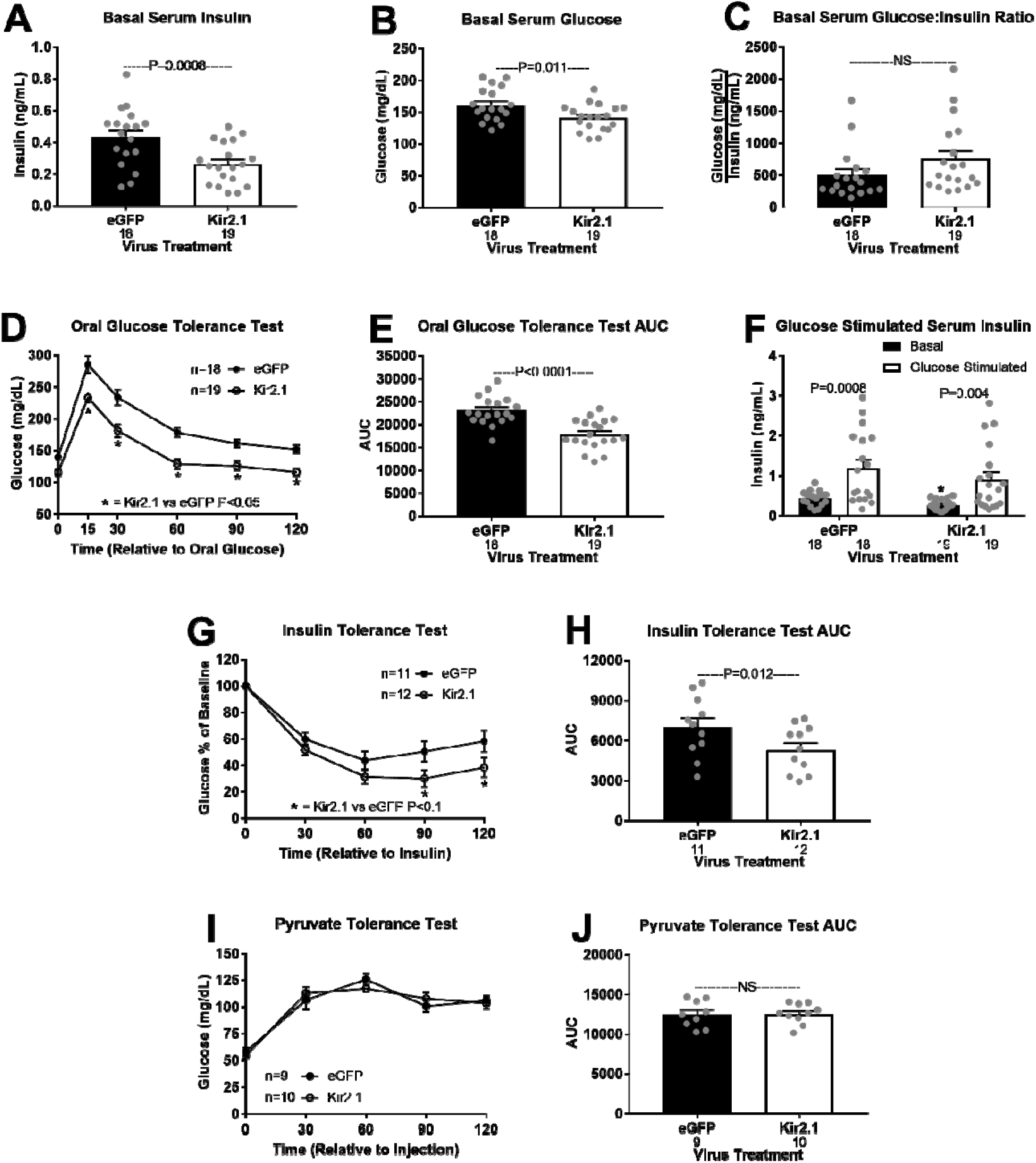
Hepatic Kir2.1 expression alters glucose homeostasis in the lean mouse. Hepatic Kir2.1 expression effects on serum insulin (A) glucose (B), glucose:insulin ratio (C), oral glucose tolerance (OGTT; D), OGTT area under the curve (AUC; E), oral glucose stimulated serum insulin (F; * denotes significance (P < 0.05) between bars of the same color), insulin tolerance (ITT; G) ITT AUC (H), pyruvate tolerance (PTT; I), and PTT AUC (J). NS = non-significant. Number below bar denotes n per group. All data are presented as mean ± SEM.

**Extended Data Figure 3.**
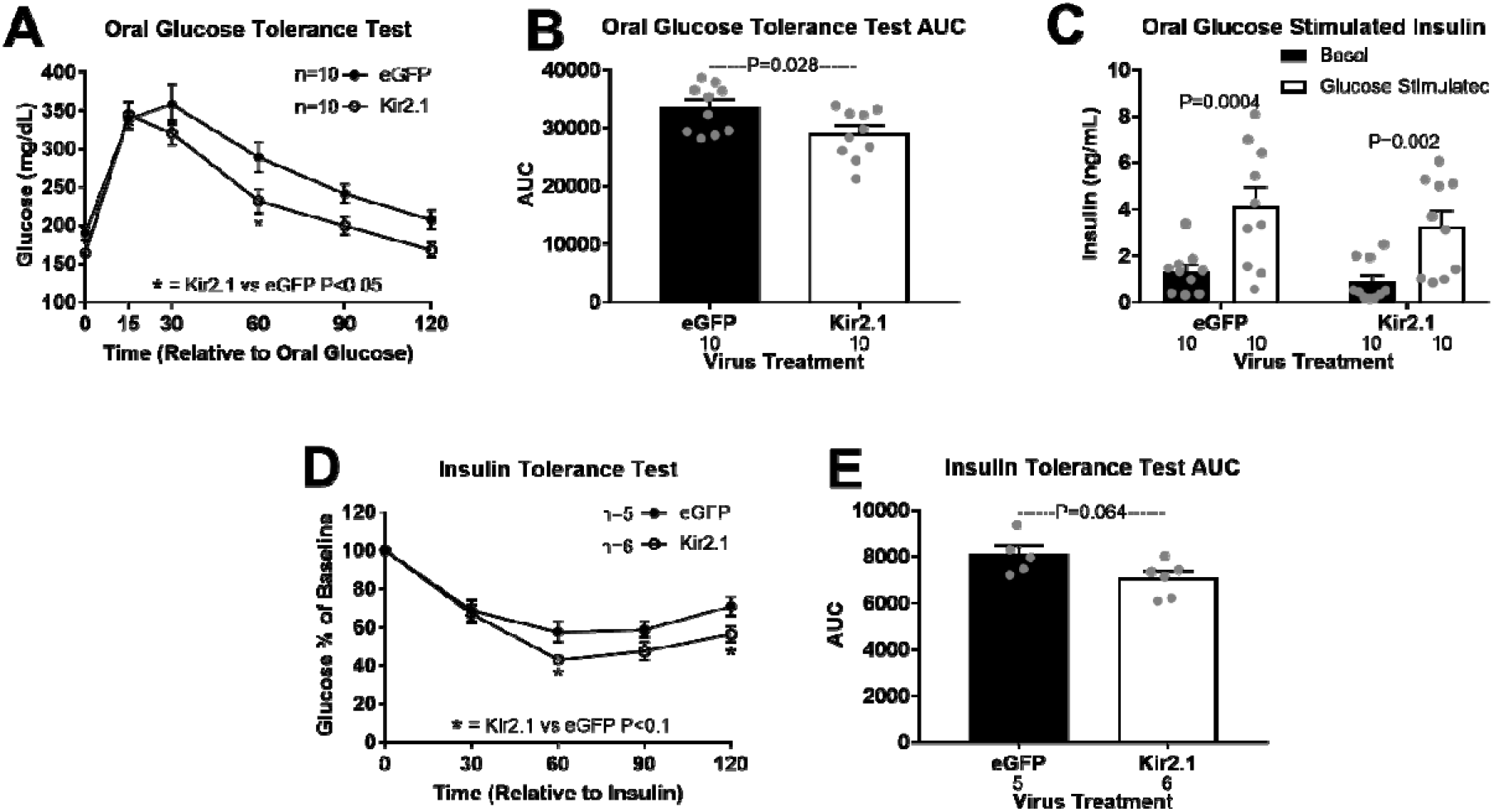
Glucose homeostasis in Kir2.1 and eGFP control mice at 3 weeks of high fat diet feeding. Effect of hepatic Kir2.1 expression on oral glucose tolerance (OGTT; A), OGTT area under the curve (AUC; B), oral glucose stimulated serum insulin (C), insulin tolerance (ITT; D), and ITT AUC (E). NS = non-significant. Number below bar denotes n per group. All data are presented as mean ± SEM.

**Extended Data Figure 4.**
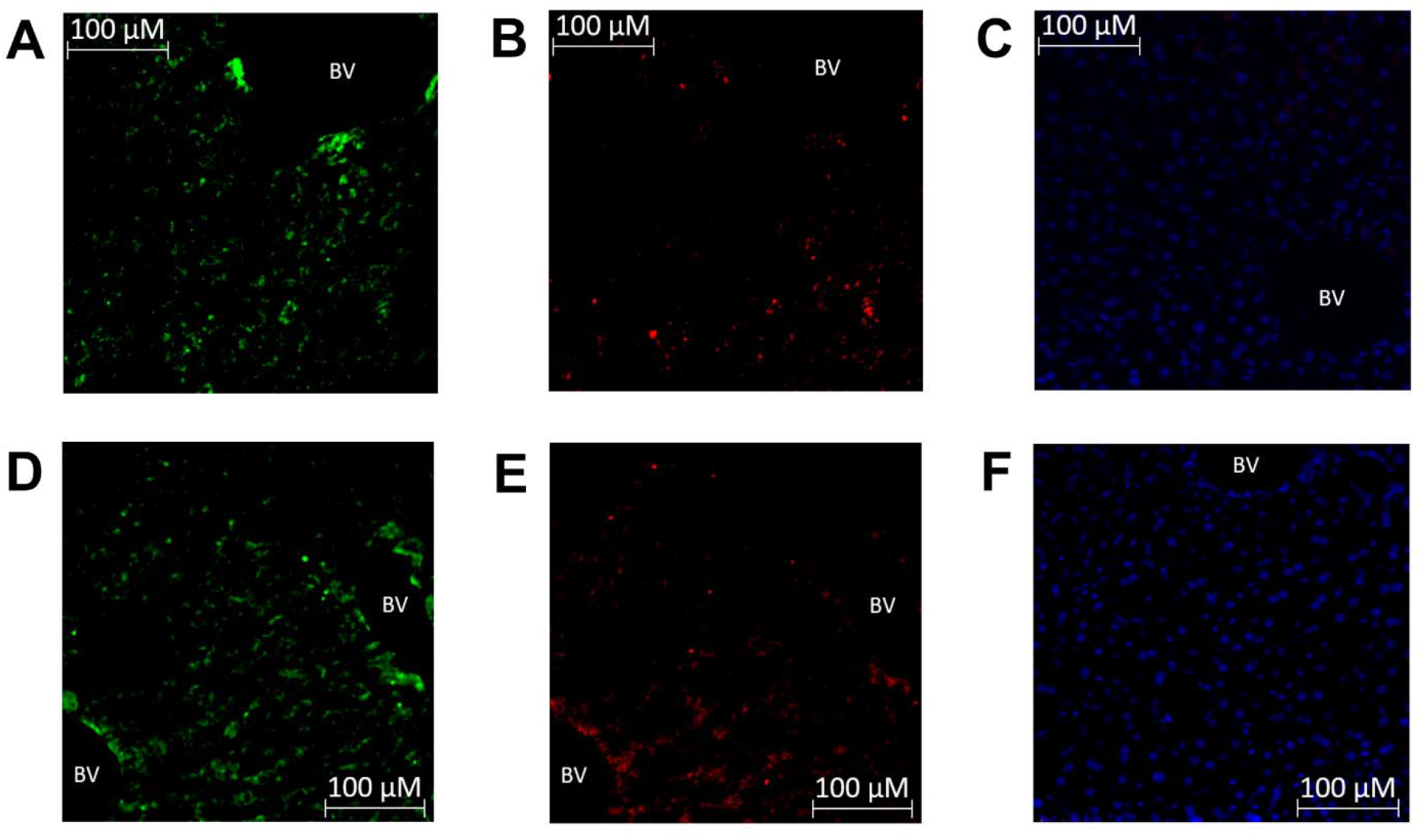
Immunohistochemical evidence of GABA_A_ receptor expressing vagal afferent innervation in the liver. Staining for the vagal afferent marker calretinin (A) and GABA_A_ receptors (B) which correspond with the co-labeled image in Fig. 6A. No primary control imaged at the same settings as Fig. 6A (C). Staining for the alternative vagal afferent marker calcitonin gene-related peptide (CGRP; D) and GABA_A_ receptors (E) which correspond with the co-labeled image in Fig. 6C. No primary control imaged at the same settings as Fig. 6C (F). Blue = DAPI (nucleus). Images at 10X magnification. BV = blood vessel

**Extended Data Figure 5.**
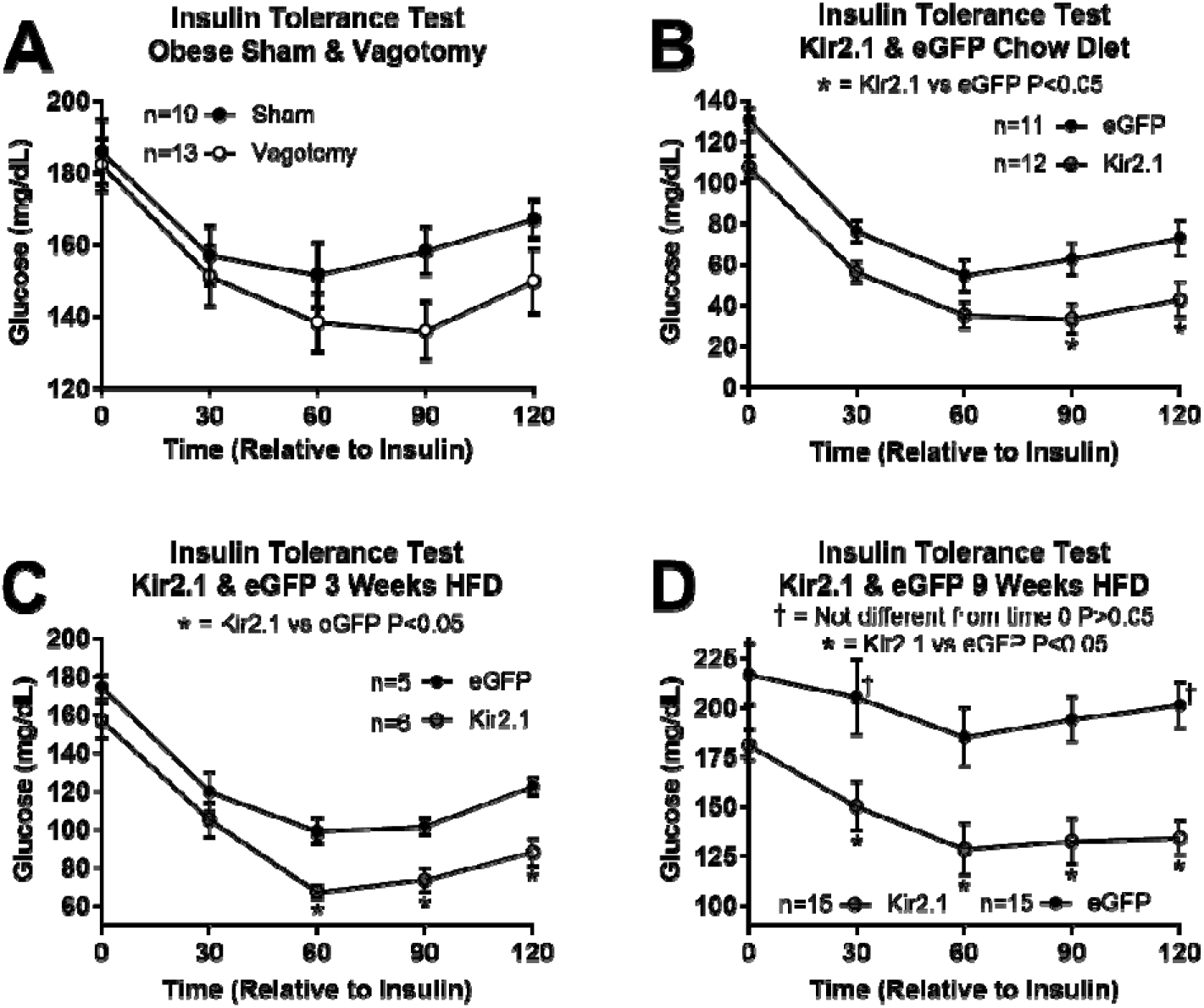
Insulin tolerance tests (ITT) presented as raw glucose values. ITT in HFD fed sham and vagotomized mice (A). ITT in Kir2.1 and eGFP control mice on chow diet (B), and after 3 (C), and 9 weeks of HFD feeding (D). † Denotes the data point is not significantly different from time 0 for that group (P > 0.05). Unless indicated, all other timepoints are significantly different from time 0 within a group of mice. * Denotes significance between groups specified in the panel within a timepoint. All data are presented as mean ± SEM.

**Extended Data Table 1.**
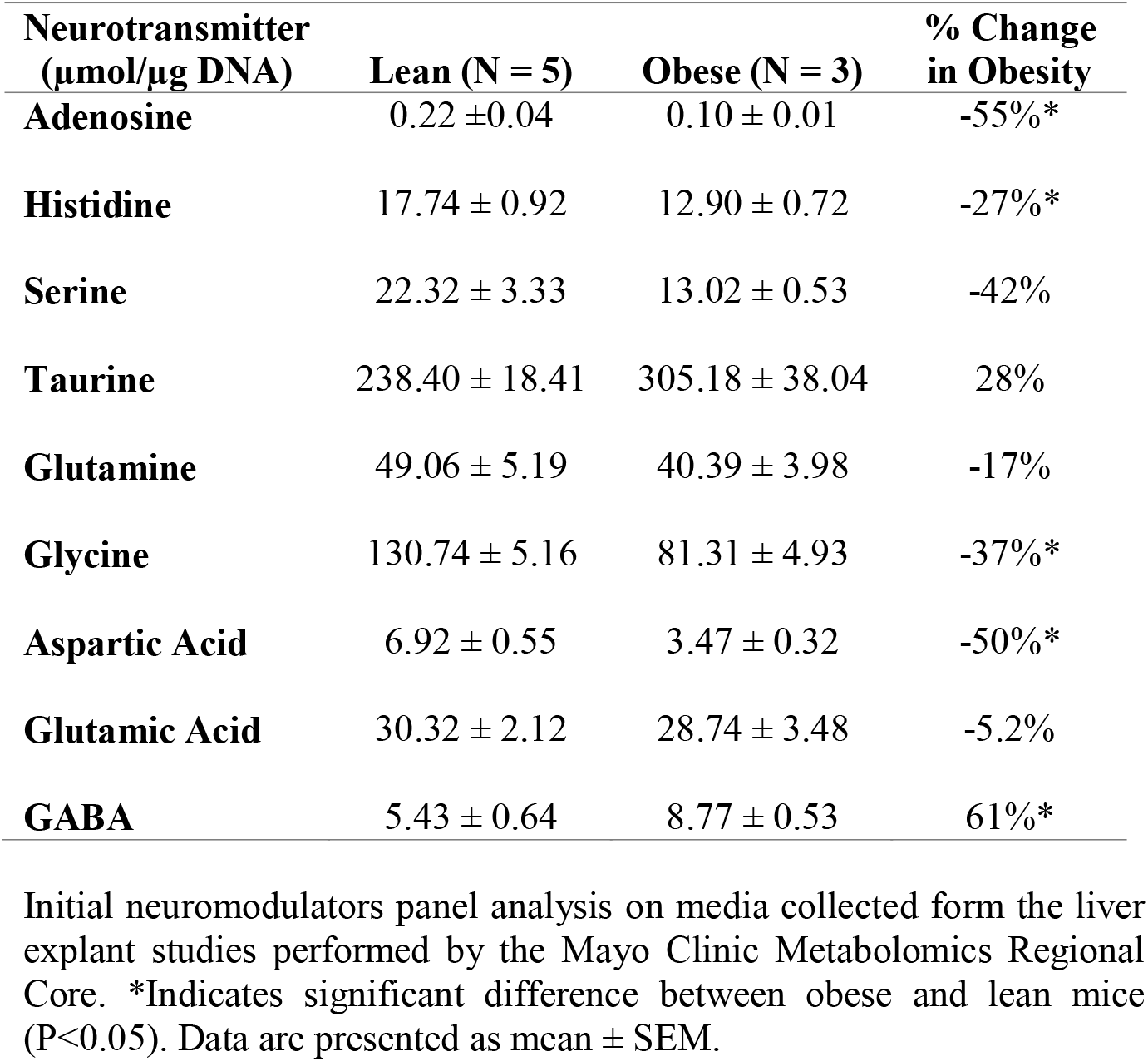
Liver slice neurotransmitter panel data

| Neurotransmitter<br>($\mu\text{mol}/\mu\text{g DNA}$) | Lean (N = 5) | Obese (N = 3) | % Change<br>in Obesity |
| --- | --- | --- | --- |
| Adenosine | 0.22 ± 0.04 | 0.10 ± 0.01 | -55%* |
| Histidine | 17.74 ± 0.92 | 12.90 ± 0.72 | -27%* |
| Serine | 22.32 ± 3.33 | 13.02 ± 0.53 | -42% |
| Taurine | 238.40 ± 18.41 | 305.18 ± 38.04 | 28% |
| Glutamine | 49.06 ± 5.19 | 40.39 ± 3.98 | -17% |
| Glycine | 130.74 ± 5.16 | 81.31 ± 4.93 | -37%* |
| Aspartic Acid | 6.92 ± 0.55 | 3.47 ± 0.32 | -50%* |
| Glutamic Acid | 30.32 ± 2.12 | 28.74 ± 3.48 | -5.2% |
| GABA | 5.43 ± 0.64 | 8.77 ± 0.53 | 61%* |

Initial neuromodulators panel analysis on media collected from the liver explant studies performed by the Mayo Clinic Metabolomics Regional Core. \*Indicates significant difference between obese and lean mice (P<0.05). Data are presented as mean ± SEM.

**Extended Data Table 2.** Metabolic characteristics of the study subjects (n=19). FFM, fat free mass; Glucose Rd, glucose disposal rate.

| | Mean ± SEM | Range |
| --- | --- | --- |
| Body mass index (kg/m<sup>2</sup>) | 45.1 ± 1.3 | 35.9 - 55.6 |
| Intrahepatic triglyceride content (%) | 11.4 ± 1.9 | 2.7 - 28.0 |
| Glucose (mg/dL) | 97 ± 2 | 81 - 121 |
| Insulin (μU/mL) | 24.1 ± 1.7 | 13.1 - 46.5 |
| Glucose infusion rate during insulin infusion (μmol/kg FFM/min) | 36.0 ± 3.0 | 15.2 - 60.8 |
| Glucose Rd during insulin infusion (% increase) | 131 ± 19 | 30 - 355 |

**Extended Data Table 3.** Regression coefficient estimates showing the association between hepatic mRNA expression of genes involved in GABA production (ABAT) and GABA transport (Slc6A6, A8, A12, and A13) and glucose infusion rate (μMol/Kg Fat Free Mass/min) and Glucose Rd (rate of disposal; % increase) during a hyperinsulinemic-euglycemic clamp.

| Glucose Infusion Rate ($\mu\text{Mol/Kg}$ Fat Free Mass/min) | | | | | |
| --- | --- | --- | --- | --- | --- |
|  | Estimate | SEM | Lower CI | Upper CI | P- Value |
| Intercept | -36.41 | 53.75 | -158.00 | 85.18 | 0.5152 |
| IHTG (%) | -0.50 | 0.19 | -0.92 | -0.08 | 0.0242 |
| SLC6A12 | 13.80 | 5.72 | 0.86 | 26.74 | 0.0391 |
| SLC6A13 | 10.74 | 4.18 | 1.28 | 20.20 | 0.0302 |
| SLC6A6 | -15.63 | 3.20 | -22.87 | -8.38 | 0.0009 |
| SLC6A8 | -5.65 | 1.95 | -10.07 | -1.23 | 0.0179 |
| ABAT | -3.26 | 7.04 | -19.20 | 12.67 | 0.6545 |
| Glucose Rd During Insulin Infusion (% Increase) |  |  |  |  |  |
|  | Estimate | SEM | Lower CI | Upper CI | P- Value |
| Intercept | -3.91 | 4.24 | -13.49 | 5.68 | 0.3805 |
| IHTG (%) | -0.03 | 0.01 | -0.07 | 0.00 | 0.0427 |
| SLC6A12 | 1.02 | 0.45 | 0.00 | 2.04 | 0.0505 |
| SLC6A13 | 0.64 | 0.33 | -0.10 | 1.39 | 0.0834 |
| SLC6A6 | -0.71 | 0.25 | -1.28 | -0.14 | 0.0204 |
| SLC6A8 | -0.45 | 0.15 | -0.79 | -0.10 | 0.0178 |
| ABAT | -0.24 | 0.56 | -1.50 | 1.01 | 0.6712 |

**Extended Data Table 4.** Single nucleotide polymorphisms (SNPs) that result in missense mutations in GABA transporters are associated with an increased incidence (OR; odds ratio) of type 2 diabetes (T2D; source: knowledge portal diabetes database). MAF – minor allele frequency.

| *SLC6A12* - T2D Associated SNPs |  |  |  |  |  |  |  |
| --- | --- | --- | --- | --- | --- | --- | --- |
| Variant ID | dbSNP ID | Predicted Impact | Study | P-value | Effect | OR | MAF |
| 12_313839_G_A | rs188610 | Missense: synonymous variant | AMP T2D-GENES T2D exome sequence analysis | 0.0238 | ↑ | 1.1 | 0.0386 |
| 12_313824_G_A | rs199521597 | Missense: early stop codon | BioMe AMP T2D GWAS | 0.0409 | ↑ | 15.8 | 0.000269 |
| *SLC6A13* - T2D Associated SNPs |  |  |  |  |  |  |  |
| Variant ID | dbSNP ID | Predicted Impact | Study | P-value | Effect | OR | MAF |
| 12_330193_C_T | rs61741313 | Missense: Replaces Arginine with Glutamine | DIAMANTE (European) T2D GWAS | 0.04 | ↑ | 1.04 | 0.01291-0.0531 |
| *SLC6A6* -T2D Associated SNPs |  |  |  |  |  |  |  |
| Variant ID | dbSNP ID | Predicted Impact | Study | P-value | Effect | OR | MAF |
| 3_14489107_G_A | rs62233560 | Missense: Replaces Valine with Isoleucine | AMP T2D-GENES T2D exome sequence analysis: Europeans | 0.00143 | ↑ | 1.4 | 0.005-0.0165 |
| 3_14523296_G_A | rs41284017 | Missense: Replaces Valine with Isoleucine | 70KforT2D GWAS | 0.00234 | ↑ | 1.38 | 0.0062-0.0165 |
| *SLC6A12* - T2D Adjusted for BMI Associated SNPs |  |  |  |  |  |  |  |
| Variant ID | dbSNP ID | Predicted Impact | Study | P-value | Effect | OR | MAF |
| 12_302492_C_G | rs138178078 | Missense: Replace Tryptophan with Serine | ExTexT2D exome chip analysis | 0.000071 | ↑ | 1.26 | 0.0052 |
| 12_319125_A_G | rs557881 | Missense: Replace Cysteine with Arginine | ExTexT2D exome chip analysis | 0.0396 | ↑ | 1.01 | 0.48 |
| 12_300248_C_G,T | rs147574089 | Missense: Replace Glutamate with Glutamine | CAMP GWAS | 0.0397 | ↑ | 6.09 | 0.0012 |
| *SLC6A13* - T2D Associated SNPs |  |  |  |  |  |  |  |
| Variant ID | dbSNP ID | Predicted Impact | Study | P-value | Effect | OR | MAF |
| 12_346454_C_T | rs140951084 | Missense in Splice Region: Replace Arginine with Glutamine | BioMe AMP T2D GWAS | 0.0359 | ↑ | 3.55 | 0.0019 |

### Materials and Methods

#### Animals

All studies excluding those done in albumin-cre expressing mice were conducted using male wildtype C57BL/6J purchased from Jackson Laboratories or bred in-house (Bar Harbor, ME). Albumin-cre male mice were purchased from Jackson Laboratories and crossed with wildtype females to generate in-house breeding of experimental albumin-cre expressing mice (Alb^Cre/+^) and sibling wildtype mice (Alb^+/+^). Mice were kept on a 14-hour light/10-hour dark schedule and housed 3-5 mice per cage until 1 week prior to study initiation, at which point animals were individually housed. We conducted studies in lean chow fed mice (7013 NIH-31, Teklad WI, 3.1 kcal/g, 18% kcal from fat, 59% kcal from carbohydrate, 23% kcal from protein) at 12-16 weeks of age. Studies in diet-induced obese sham and vagotomy mice were performed after 9 weeks on a high fat diet (TD 06414, Teklad WI, 5.1 kcal/g, 60.3% kcal from fat, 21.3% kcal from carbohydrate, 18.4% kcal from protein; 20-26 weeks of age). Studies in obese Kir2.1 expressing mice were performed at 3, 6, and 9 weeks after introduction of the high fat diet and all studies were repeated in 3 different cohorts. Kir2.1 and eGFP expressing mice weighing under 36 grams after 9 weeks of high fat diet feeding were excluded from all data. Unless fasted, mice had ad libitum access to food and water. All studies were approved by the University of Arizona Institutional Animal Care and Use Committee.

#### Hepatic Vagotomy Surgeries

Surgeries were performed in 12-week old male C57BL/6J mice under isoflurane anesthesia. Mice were randomly assigned to a surgical group (sham or vagotomy). A ventral midline incision through the skin and peritoneum allowed us to isolate the hepatic vagus nerve as it branched from the esophagus (Fig. 1A). In vagotomized mice, we severed the hepatic vagal nerve (Fig. 1A; arrow A), while it remained intact in sham operated mice. The peritoneum was sutured with absorbable polyglactin 910 suture and the skin with nylon suture. Mice were given a single post-operative dose of slow release formulated buprenorphine analgesic (1.2 mg/kg slow release, sub-cutaneous). We monitored food intake and body weight daily and removed sutures 7 days post-operation.

#### Viral Induced Channel Expression

The depolarizing channel (PSAM^L141F,Y115F^-5HT3HC), originally engineered by Dr. Scott Sternsons group ^14^, was made by mutating the acetylcholine binding domain of a chimeric channel that included the binding domain of the α7 nicotinic acetylcholine receptor and the ion pore domain of the serotonin receptor 3a. The ligand binding domain mutations (Leu^141^ → Phe and Tyr^115^ → Phe) limited the agonist action of acetylcholine and allowed for stimulation by a pharmacologically selective effector molecule PSEM89S. The exogenous ligand PSEM89S opens the serotonin receptor 3a channel allowing Na^+^, K^+^, and Ca^++^ passage into the cell and membrane depolarization. AAV8 viral vectors were used for plasmid delivery in all the reported studies and were synthesized by the Penn Vector Core. Hepatic specific expression of the depolarizing channel was achieved through two different methods. First, expression of a cre-recombinase dependent depolarizing channel was driven by a globally expressed CAG promoter. LoxP sites limited expression to cre-recombinase expressing tissue, and tail vein injection of 1×10^10^ viral genome copies established hepatocyte expression in albumin-cre but not wildtype mice. Second, a separate AAV8 viral vector, induced hepatic specific expression of the same depolarizing channel by driving expression using the thyroxine binding globulin (TBG) promoter. Tail vein injection of 1×10^11^ viral genome copies established hepatocyte expression (Fig. 2C and Extended Data Fig. 1B). The thyroxine binding globulin promoter also drove hepatic expression of the hyperpolarizing, inward-rectifier K^+^ channel, Kir2.1. Tail vein injection of 1×10^11^ viral genome copies established hepatocyte specific expression (Fig. 3A). To confirm channel expression and tissue specificity, all viral vector plasmids encoded for enhanced green fluorescent protein (eGFP).

#### PSEM89S Ligand Injection Studies

All studies in virus injected mice were conducted at least 5 days post virus injection to allow for maximal channel expression. Individually housed mice were intraperitoneally injected with the ligand for the depolarizing channel (PSEM89S; 30mg/kg; 0.1mL/10g body weight) or PBS (0.1mL/10g body weight).

Studies conducted in mice injected with the cre-dependent depolarizing channel virus took place at 8 am. Food was removed upon study initiation. Blood for serum insulin and glucose determination was collected from the tail vein 15 minutes following intraperitoneal injection. All mice received both saline and PSEM89S ligand injection on separate days.

Studies conducted in mice expressing the cre-independent depolarizing channel began at 1 pm following a 4 hour fast. Mice received an oral glucose gavage (2.5 g/kg) 10 minutes after intraperitoneal injection of the PSEM89S ligand or saline. 15 minutes following glucose administration (25 minutes post treatment injection), blood for serum insulin and glucose determination was collected from the tail vein. All mice received both saline and PSEM89S ligand injection on separate days. These studies were repeated in 2 cohorts.

#### Electrophysiology

We performed simultaneous in vivo recordings of hepatocyte membrane potential and hepatic afferent vagal nerve firing activity in anesthetized (isoflurane) mice to directly assess the effect of hepatocyte depolarization on hepatic afferent vagal nerve activity. The abdomen was shaved and scrubbed with betadine and isopropanol before an incision through the skin and peritoneum was made to expose the internal organs. The intestines were moved to expose the liver and one lobe of the liver was secured onto a small platform to minimize movement caused by respiration. A ground electrode was secured under the skin and the hepatic vagal nerve was gently lifted onto a hook-shaped electrode (Fig. 1A; arrow A) attached to the positive pole of a Grass P511 AC coupled amplifier, and the signal was filtered with a bandwidth of 300-1000 Hz. The nerve and hook electrode were dried and surrounded with ice cold kwik-sil to secure placement of the hook. The hepatic vagal nerve to the right of the hook, near the esophagus was cut to eliminate efferent firing (Fig. 1A; arrow B). Once the kwik-sil had set, the anesthetized mouse was bathed in 37◻C Krebs-Henseleit (KH) buffer gassed with CO2. After placement and sealing of the hook electrode, 45-60 minutes of basal nerve activity was monitored/recorded with pClamp software (version 10.2; Molecular Devices) until nerve activity stabilized, after which, we began treatments.

Simultaneously, intracellular recordings of hepatocytes were made with sharp glass electrodes (30-40 MΩ) pulled from thin-walled borosilicate glass capillary tubes (OD: 1 mm; ID: 0.78mm; Sutter Instrument Co., Novato, CA), filled with 1.5M KCl and positioned visually using a motorized 4-axis micromanipulator (Siskiyou, Grants Pass, OR). Electrical signals were conducted via an Ag–AgCl electrode connected to a headstage (Axoclamp ME-1 probe), which was in turn connected to an Axoclamp 2B amplifier. Both nerve and intracellular signals were sent to an A/D converter (Digidata 1322A, Molecular Devices, Sunnyvale, CA), digitized at 20 kHz and viewed on a computer monitor using pClamp software (version 10.2; Molecular Devices).

Before treatments were applied hepatocyte impalement was determined by an abrupt negative deflection upon penetration of the cell and a stable intracellular potential (−45 to −25 mV for mouse hepatocytes) for at least 2 minutes. If the recording of hepatocyte membrane potential was not stable the electrode was removed and membrane potential was measured on another hepatocyte.

To assess the response to channel activation, PSEM89S ligand was bath applied (30 μM) for 45 min during recordings. In order to understand the effect of Kir2.1 channel on hepatocyte membrane potential, a 10-minute baseline measure was collected and then Barium (BaCl 50 μM) was bath applied and recording continued for 45 minutes. Barium blocks Kir2.1 mediated current, thus the change in membrane potential in response to barium indicates the degree of hyperpolarization resulting from Kir2.1 channel expression. In all electrophysiology studies mice were sacrificed by cutting the diaphragm and subsequent cervical dislocation. Tissues were collected to confirm tissue specificity of channel expression and the % of hepatocytes that were expressing the channel. All studies were performed at room temperature (25°C).

#### Immunohistochemistry and Imaging

To confirm the specificity and extent of viral-induced channel expression in hepatocytes, immunohistochemistry for GFP was performed. Liver, adipose, pancreas, and skeletal muscle were collected into 4% paraformaldehyde in 0.1 M PBS (Phosphate Buffered Saline) immediately after sacrifice. After 4 h at 4°C, tissues were transferred to a 30% sucrose solution in 0.1 M PBS and kept at 4°C until tissues sunk to the bottom of the solution. Tissues were snap frozen on liquid nitrogen in OTC (Optimal Cutting Temperature; Sakura Finetek USA Inc, Torrance, CA) and stored at −80°C. We used a cryostat HM 520 (MICROM International GmbH, Walldorf, Germany) to get 10 μM thick slices which we collected onto Superfrost Plus slides. Immunohistochemistry for GFP alone (Figs. 2A-2C and Extended Data Figs. 1A-1B) was performed as follows: Briefly, slides were washed twice in PBS and twice in PBST (3% Triton in PBS) before being exposed to blocking solution (5% normal goat serum in PBST) for 1 h. Slides were subsequently exposed to a 1:5000 dilution of the primary anti-GFP antibody in blocking solution (Alexa488-conjugated rabbit anti-GFP; Life Technologies, Waltham, MA) for 3 hours at room temperature. After primary antibody incubation we washed the slides 3 times in PBST and 2 times in PBS prior to placing the coverslip with DAPI Fluoromount-G as the mounting medium (SouthernBiotech, Birmingham, AL). Fluorescent imaging was performed without antibody amplification in mice administered the AAV8 that encoded for Kir2.1 and tdTomato. Immunohistochemistry for GFP and the hepatocyte specific marker arginase-1 (Extended Data Figs. 1C-1E) was performed as follows: Slides were washed twice in PBS and twice in PBST (3% Triton in PBS) before being exposed to blocking solution (5% normal donkey serum in PBST) for 1 h. Slides were then incubated overnight at 4°C in a 1:400 dilution of the primary anti-arginase-1 antibody in blocking solution (Rabbit anti-liver arginase ab91279; Abcam, Cambridge, UK). After overnight primary antibody incubation slides were washed 5 times with PBST and exposed to a 1:500 dilution of the primary anti-GFP antibody (Alexa488 conjugated mouse anti-GFP sc-9996 AF488; Santa Cruz, Dallas, TX) and the secondary anti-rabbit antibody (Alexa568 conjugated donkey anti-rabbit A10042; Thermo Fisher, Waltham, MA) in blocking solution and for 1 hour at room temperature. Slides were then washed 5 times in PBST and 2 times in

PBS prior to applying DAPI and a coverslip. Immunohistochemistry for calcitonin gene-related peptide (CGRP) and GABA_A,_ and calretinin and GABA_A_ were performed identical to that described for GFP and arginase-1 with the following primary anti-CGRP and anti-GABA_A_ antibodies at a 1:100 dilution (Goat anti-CGRP ab36001 and rabbit anti-GABA_A_ α5 ab10098; Abcam, Cambridge, UK) and the primary anti-calretinin antibody at a final concentration of 15 μg/mL (Goat anti-calretinin AF5065; R&D Systems, Inc., Minneapolis, MN). The secondary anti-rabbit and anti-goat antibodies were used at a 1:500 dilution (Alexa568 conjugated donkey anti-rabbit A10042 and alexa488 conjugated donkey anti-goat A32814; Thermo Fisher, Waltham, MA). Images were collected by fluorescent microscopy (Leica DM5500B, Leica Microsystems, Wetzlar, Germany), captured using HCImage Live, and formatted in Image-Pro Premier 9.2. 10X magnification was used to ensure a wide field of vision and accurate assessment of degree of expression. 20X magnification was used to image co-staining for GFP and arginase-1.

#### Glucose Tolerance Test

Oral glucose (2.5g/kg; 0.1mL/10g body weight; Chem-Impex Int’l Inc., Wood Dale, IL) was given to 4 hour fasted individually housed mice. All glucose tolerance tests began at 1 pm and glucose was measured in whole blood, collected from the tail vein, by glucometer (Manufacture # D2ASCCONKIT, Bayer, Leverkusen, Germany) at 0, 15, 30, 60, 90, and 120 minutes after glucose gavage. Blood for serum insulin (oral glucose stimulated insulin secretion; OGSIS) and glucose determination was collected from the tail vein 15 minutes following glucose administration.

#### Insulin Tolerance Test

Intraperitoneal insulin (0.75U/kg; 0.1mL/10g body weight; Sigma Aldrich, St. Louis, MO) was given to 4 hour fasted individually housed mice. All insulin tolerance tests began at 1 pm and glucose was measured in whole blood, collected from the tail vein, by glucometer (Manufacture # D2ASCCONKIT, Bayer, Leverkusen, Germany) at 0, 30, 60, 90, and 120 minutes after insulin injection.

#### Pyruvate Tolerance Test

Intraperitoneal sodium pyruvate (1.5g/kg; 0.1mL/10g body weight; Alfa Aesar, Ward Hill, MA) was given to 16 hour fasted individually housed mice. Mice were switched to wood chip bedding (Harlan Laboratories; Cat. # 7090 Sani-Chips) at the initiation of the fast. All pyruvate tolerance tests began at 9 am and the rise in glucose was measured in whole blood, collected from the tail vein, by glucometer (Manufacture # D2ASCCONKIT, Bayer, Leverkusen, Germany) at 0, 30, 60, 90, and 120 minutes after pyruvate injection. This is indicative of hepatic gluconeogenic potential from pyruvate.

#### Serum Assays

Within 2 hours of collection, blood was left to clot at room temperature for 20 minutes. Thereafter the blood was centrifuged at 3,000xg for 30 minutes at 4°C and serum was collected. Serum was stored at −80°C until metabolite and hormone analyses. Serum glucose was analyzed by colorimetric assay (Cat. # G7519, Pointe Scientific Inc., Canton MI). Serum insulin was analyzed by enzyme-linked immunosorbent assay (ELISA; Cat. # 80-INSMSU-E01,E10, Alpco, Salem, NH). Serum glucagon was analyzed by enzyme-linked immunosorbent assay (ELISA; Cat. # 10-1281-01, Mercodia, Uppsala, Sweden) from tail vein blood collected at 9 am from fed mice (Vagotomy study) or after a 4 hour fast at 1 PM (Kir2.1 study).

#### Liver Slice Explant Studies

Liver slices from experimental mice were incubated *ex vivo* to measure release of signaling molecules. A peristaltic pump perfusion system was used to deliver warmed KH buffer to the liver through the portal vein. Briefly, mice were anesthetized with an intraperitoneal injection of ketamine (10mg/mL) and diazepam (0.5mg/mL). Once mice were unresponsive, an incision in the lower abdomen through the skin and peritoneal membrane was made vertically through the chest along with transverse incisions on both sides to expose the liver. A 30-gauge needle was inserted into the hepatoportal vein to blanch the liver. The inferior vena cava was cut to relieve pressure in the circulatory system and allow blood to drain. The perfusion continued for several minutes at a rate of 4mL/minute until the liver was completely blanched. The liver was removed and washed in warm PBS before being sliced into 0.2 mm slices using a Thomas Sadie-Riggs Tissue Slicer. Two liver slices were taken from each mouse. Tissue slices were placed individually into a well on a 12-well plate pre-filled with 1mL of KH buffer that had been sitting in an incubator set to 37°C and gassed with 5% CO_2_. Liver slices were incubated in the initial well for 1 hour to stabilize before being transferred to a fresh well pre-filled with KH buffer. Liver slices treated with the GABA-T inhibitor EOS were incubated in media containing EOS (5.3 mM) during the second hour of incubation. Liver slices treated with the Na^+^/K^+^ ATPase inhibitor ouabain were incubated in media containing ouabain (1 mM) media was collected after 15 minutes of incubation. Liver slices treated with the GABA transporter inhibitors for BGT1 and GAT2 were incubated in media containing betaine (1 mM) or Nipecotic acid (1 mM) or both during the second hour of incubation. Liver slices incubated in reduced and low NaCl media sat in normal KH buffer (NaCl 118 mM) for the first hour and then were transferred to reduced (60 mM) or low NaCl (15 mM) KH buffers for the second hour. For the reduced and low NaCl medias, respectively, 58 and 103 mM of NaCl were replaced with 116 and 206 mM of mannitol to maintain the osmolarity of the buffer. After 1 hour in the second well, tissue and media were collected. Liver slice samples and KH media samples from both wells of each mouse were pooled. Liver slices were snap frozen in liquid nitrogen, while media was frozen and stored at −80°C for future analysis.

#### Explant Media Analysis

Preliminary media samples were sent to the Mayo Clinic Metabolomics Regional Core for mass spectrophotometry analysis using their neuromodulators panel (Extended Data Table 1). For all liver slice GABA and aspartate release data, we thawed the media collected from the *ex vivo* hepatic slice culture on ice and centrifuged for 5 minutes at 10,000xg at 4°C to remove tissue debris. We then measured GABA in the supernatant using a commercially available ELISA (REF# BA E-2500, Labor Diagnostika Nord, Nordhorn, Germany).

Aspartate release was measured using liquid chromatography-mass spectrometry. Samples were prepared for analysis by LC-MS/MS using protein precipitation. Twenty μl of each sample and standard curve increment was transferred to 1.5 ml tubes. One hundred eighty μl acetonitrile (ACN) was added to each tube followed by a 5 second vortex. All samples were incubated at 4°C for one hour for precipitation. Samples were then centrifuged at 10,000 RPM for 10 minutes and the supernatant transferred to 300 μl HPLC vials for analysis. The aqueous portion of the mobile phase was buffered using 10 mM ammonium bicarbonate with the pH adjusted to 7.4 using 1M formic acid and ammonium hydroxide. Methanol was used as the organic portion of the mobile phase. The column for separation was a Phenomenex Luna Silica(2) with 5 μm particle diameter and 100 Å pore size. Column internal diameter was 4.6 mm and length was 150 mm. A Shimadzu LC10 series HPLC with two dual piston pumps was used for sample injection and solvent delivery. The flow rate was fixed at 300 μl per minute. Aspartate was quantified using an LTQ Velos Pro mass spectrometer. Eluate from the Shimadzu HPLC was ionized using a Thermo ESI source. Source voltage was 6 kV; sheath and auxiliary gas flows were 40 and 20 units respectively. The ion transfer capillary was heated to 300°C. The LTQ Velos Pro was operated in negative SRM mode using two transitions: 132.1->115 for quantification and 132.1->88.1 as a qualifier. Data integration and quantification were performed using the Thermo Xcalibur software packaged with the LTQ Velos Pro.

#### Liver Analyses

Prior to analysis, frozen livers were powdered using a liquid nitrogen cooled mortar and pestle to obtain homogenous liver samples. To measure liver DNA content (ng dsDNA/g tissue), 10-20 mg of powdered liver was sonicated in 200μL DEPC H_2_O and dsDNA determined by fluorometric assay (Cat. # P7589, Invitrogen, Waltham, MA). Whole liver mRNA was isolated from powered liver samples with TRI Reagent® (Life Technologies, Grand Island, NY) and phenol contamination was eliminated by using water-saturated butanol and ether as previously described ^51^. cDNA was synthesized by reverse transcription with Verso cDNA synthesis kit (Thermo Scientific, Inc., Waltham, MA), and qPCR performed using SYBR 2X mastermix (Bio-Rad Laboratories, Hercules, CA) and the Biorad iQ™5 iCycler (Bio-Rad Laboratories, Hercules, CA). Expression of β-actin (ACTβ) and GABA-Transaminase (ABAT) mRNA were measured using the following primers (5’→3’): ACTβ; Forward – TCGGTGACATCAAAGAGAAG Reverse – GATGCCACAGGATTCCATA, ABAT; Forward – CTGAACACAATCCAGAATGCAGA Reverse - GGTTGTAACCTATGGGCACAG. LinReg PCR analysis software was used to determine the efficiency of amplification from raw output data ^52^. ACTβ served as the reference gene for calculating fold change in gene expression using the efficiency^−ΔΔCt^ method ^53^.

Total liver lipids were extracted from powdered liver samples. Briefly, 10-20 mg of sample was sonicated in 100μL PBS. 1 mL of 100% ethanol was added to each sample and vortexed for 10 minutes. Following 5 minutes of centrifugation at 16,000xg at 4°C, supernatant was transferred to a fresh tube for analysis of liver triglycerides (Cat. # T7531, Pointe Scientific Inc., Canton, MI).

Hepatic NADH and NAD were quantified by fluorometric assay (ab176723, Abcam, Cambridge, UK). Hepatic ATP concentrations were assessed as previously described ^54^.

#### Studies Conducted in Human Subjects

A total of 19 men and women with obesity who were scheduled for bariatric surgery at Barnes-Jewish Hospital in St. Louis, MO participated in this study, which was conducted at Washington University School of Medicine in St. Louis. Subjects provided written, informed consent before participating in this study, which was approved by the Human Research Protection Office at Washington University School of Medicine in St. Louis, MO (ClinicalTrials.gov NCT00981500). Intrahepatic triglyceride content was determined by using magnetic resonance imaging (3.0-T superconducting magnet; Siemens, Iselin, NJ) in the Center for Clinical Imaging Research. A 7-hour (3.5-h basal period and 3.5-h insulin infusion period) HECP, in conjunction with stable isotopically labelled glucose tracer infusion, was then conducted in the Clinical Translational Research Unit (CTRU), as previously described^55^. This procedure was performed to determine whole-body insulin sensitivity, which was assessed as the glucose infusion rate (expressed as mg·kg FFM^−1^·min^−1^) during during the last 30 minutes of the HECP; and skeletal muscle insulin sensitivity, which was assessed as the percent increase in the rate of glucose disposal above basal, during the last 30 minutes of the HECP ^55^.

#### Liver RNA sequencing (RNA-seq)

Liver tissue was obtained by needle biopsy during the bariatric surgical procedure, before any intra-operative procedures were performed. Liver tissue was rinsed in sterile saline, immediately frozen in liquid nitrogen, then stored at −80°C until RNA extraction. Total RNA was isolated from frozen hepatic tissue samples by using Trizol reagent (Invitrogen, Carlsbad, CA). Library preparation was performed with total RNA and cDNA fragments were sequenced on an Illumina HiSeq-4000. The fragments per kilobase million reads upper quartile (FPKM-UQ) values were calculated and used for further gene expression analyses. All RNA-seq data used in this study have been deposited into the NCBI GEO database under accession number GSE144414.

#### Statistics

We analyzed the data in SAS Enterprise Guide 7.1 (SAS Inst., Cary, NC), using a mixed model ANOVA for all analyses. ANOVA tests do not have a one-tailed vs. two-tailed option, because the distributions they are based on have only one tail. When comparisons between all means were required, we used a Tukey’s adjustment for multiple comparisons. When comparisons of means were limited (e.g. only within a timepoint or treatment), we used a bonferonni correction for multiple comparisons. For the analysis of ITT, OGTT, and PTT repeated measures ANOVA were performed by including time point in the analysis. When applicable analyses were conducted separately for chow and HFD fed mice. In the Kir2.1 mice, which were monitored for response at 0, 3, 6, and 9 weeks, analyses were performed for each timepoint individually. For analysis of cre-dependent depolarizing channel effects, analysis was performed in each genotype separately (Alb^cre/+^ or Alb^+/+^) and the main effect was injection (PSEM89S ligand or saline). For analysis of the studies using the thyroxine binding promoter driven ligand gated depolarizing channel we had the main effect of injection (PSEM89S ligand or saline). For the vagotomy analyses the main effect was surgery (sham or vagotomy) and weeks on high fat diet when applicable. For the Kir2.1 analyses the main effect was virus (eGFP or Kir2.1). Linear regressions of body weight and serum insulin concentrations were performed on Kir2.1 and eGFP controls, and sham and vagotomized mice using SAS Enterprise Guide 7.1. All insulin tolerance tests are presented as a percentage of baseline glucose in main and Extended Data figures, and additionally presented as raw glucose values in Extended Data Fig. 5. Human data was analyzed using a multivariate regression including IHTG content, and *SLC6A6, SLC6A8, SLC6A12,* and *SLC6A13* mRNA expression as independent variables with Type 3 test of fixed effects used to determine significance and estimates derived from maximum likelihood estimation. All graphs were generated using GraphPad Prism 8 (GraphPad Software Inc., La Jolla, CA).

## Data Availability

The datasets generated and/or analyzed during the current study are available in the Mendeley data repository at a link provided in the cover letter. Datasets will be made public upon acceptance of this manuscript.

## Endnotes

## Acknowledgments

The authors wish to thank Dr. Scott Sternson at the Howard Hughes Medical Institute, Ashburn, VA., for providing the PSEM89S ligand used in these studies, Ken Pendarvis at the University of Arizona for analyzing aspartate release from our *ex vivo* hepatic slice culture model by liquid chromatography-mass spectrometry, The Mayo Clinic Metabolomics Regional Core for performing the initial mass spectrophotometry analysis of media from our *ex vivo* hepatic slice culture model using their neurotransmitter panel to identify potential hepatocyte released signaling molecules, Drs. Ralph Fregosi, Masoud Ghamari-Langroudi and Richard Levine for guidance on experimental models and electrophysiology.

## Author Contributions

CEG – Experimental design and project conceptualization, performed experiments and wet lab analyses, wrote initial draft of manuscript, generated figures, reviewed and edited manuscript.

CH – Performed experiments and wet lab analyses, reviewed and edited manuscript.

SG – Performed experiments and wet lab analyses, reviewed and edited manuscript.

MRH – Developed electrophysiology methodology and performed experiments, reviewed manuscript.

BJR – Experimental design and project conceptualization, performed surgeries, analyzed statistics, reviewed and edited manuscript.

## Author Information

Reprints and permissions information is available at www.nature.com/reprints. Competing Interests: The results presented in this paper have resulted in patent cooperation treaty Application No. 62/511,753 and 62/647,468: METHODS AND COMPOSITIONS FOR REGULATING GLUCOSE HOMEOSTASIS. Correspondence and requests for materials should be addressed to

